# Cardiolipin regulates free energy of folding and vesicle leakage of CM15 antimicrobial peptides in bacterial inner membranes

**DOI:** 10.64898/2026.09.05.749580

**Authors:** Nishant Mohan Bahuguna, Aditya Upasani, Avijeet Kulshrestha, Sudeep Punnathanam, Rahul Roy, Ganapathy Ayappa

## Abstract

Globally rising antimicrobial resistance has led to antimicrobial peptides (AMPs) being explored as alternative therapeutics for bacterial infections. A better mechanistic understanding of the interaction of the peptide with different membrane compositions can aid in targeting specific bacterial strains more effectively and lead to more resilient treatments. Using all-atom molecular dynamics simulations, we obtain the free-energy landscape of the cecropin–melittin hybrid peptide CM15 using a path-based method and investigate the effect of cardiolipin (CL), a four-tailed lipid found in bacterial membranes. Our analysis reveals that the transition of CM15 from an unfolded state in solution to a membrane-bound helical state proceeds through three main steps: membrane binding, insertion in an unfolded state, and subsequent folding beneath the lipid headgroups into an *α*-helical conformation. In a phosphatidylethanolamine (PE)/phosphatidylglycerol (PG) membrane lacking CL, folding occurs spontaneously without appreciable free-energy barriers. However, the addition of only 5% CL substantially alters the landscape, giving rise to two distinct transition pathways with barriers ranging from 2.5 to 8 *k*_*B*_*T* . This reflects the presence of a more rugged free-energy landscape associated with local heterogeneity in lipid composition. The emergence of these barriers is driven by strong interactions between CL and CM15, which promote CL sequestration around the peptide in regions of negative membrane curvature. Vesicle leakage experiments reveal the inhibitory influence of CL at lower peptide-to-lipid ratios with delayed kinetics as the CL content is increased. We attribute the reduced activity at low CM15 concentrations to the free-energy barriers associated with peptide folding and to strong CM15–CL interactions that stabilize a membrane bound peptide state, thereby hindering membrane disruption, pore formation, and leakage. Our study illustrates the putative role of CL in pore formation, providing molecular insights that can potentially aid the rational design of strain-specific synthetic AMPs.

**SIGNIFICANCE:** Antimicrobial peptides (AMPs) are being actively studied to combat the emergence of bacterial resistance to conventional antibiotics. AMPs act by disrupting the bacterial cell membrane; however, a molecular view of the folding transition at the membrane-water interface is incompletely understood. Our combined simulation and vesicle leakage study on CM15 provides molecular insights into the membrane insertion and folding pathways, with free-energy computations revealing the inhibitory role of the four-tailed lipid cardiolipin. We expect our results to apply across a wide class of amphipathic AMPs, assisting in the design of strain-specific peptides.

## INTRODUCTION

Antimicrobial peptides (AMPs) are an integral part of our innate immune system and have been found across different species in the animal kingdom. AMPs have been explored as potential antibacterial candidates since they have a lower propensity to induce bacterial resistance [1, 2] compared to traditional antibiotics, attributed primarily to their mechanically disruptive mode of action and co-evolution with bacteria [3]. Since bacterial membrane structures and compositions are generally more conserved, membrane-targeting AMPs offer a promising alternative therapeutic strategy to bypass bacterial resistance that arises from many conventional antibiotics [4, 5]. AMPs have been shown to have antimicrobial activity against various pathogens, including Gram-positive and Gram-negative bacteria, fungi, and viruses [6], as well as several drug-resistant bacterial strains such as ESKAPE group pathogens [7].

The cell envelope of Gram-negative strains consists of an outer lipopolysaccharide layer and an inner phospholipid bilayer with an intervening peptidoglycan layer referred to as the cell wall. Gram-positive strains are devoid of the outer lipopolysaccharide (LPS) membrane and instead have a thicker peptidoglycan cell wall with an inner membrane made up of phospholipids [8]. Despite the diverse multicomponent nature of bacterial membranes, the inner membrane of both Gram-positive and Gram-negative strains is negatively charged, rendering it a target for AMPs carrying a net positive charge. Bacteria, however, have evolved to modify their membrane charge, or zeta potential, in response to external antibacterial molecules [9].

A typical inner cell membrane of *Escherichia coli* consists of 75% 1,2-dioleoyl-sn-glycero-3-phosphoethanolamine (PE), 20% 1,2-dioleoyl-sn-glycero-3-phospho-rac-(1-glycerol) (PG), and 5% tetraoleoyl-cardiolipin (CL) [10–12]. This composition is relatively invariant under a broad spectrum of growth conditions, with a few exceptions. For instance, the CL content increases when cells enter the stationary phase [13]. In Gram-positive strains, recent studies have shown that CL content can vary substantially, from 5% in *S. aureus* to 85% in *N. lacusekhoensis*, an extremophilic strain [14]. Changes in CL content can also influence AMP activity. The AMPR-22 peptide shows lower carboxyfluorescein release and reduced activity with increasing CL content in large unilamellar vesicles [15]. An increase in CL concentration in liposomes has also been shown to increase lipid packing and reduce the activity of AMPs such as LL-37 and ΔM2 by inhibiting calcein leakage [16]. In *E. coli*, CL composition increases while PE content decreases when bacteria are subjected to osmotic stresses with electrolytes or non-electrolytes [17].

Molecular dynamics (MD) simulations have been extensively used to decipher interactions of AMPs with biological membranes [18–21]. Many amphipathic AMPs are unfolded or partially folded in solution and attain a folded state upon membrane binding and insertion. Their disruptive action can involve transmembrane pore formation as well as membrane permeabilization mechanisms that do not necessarily require stable pores [22–24]. Pore formation is a collective phenomenon involving lipid and peptide co-localization, the energetics of which can be strongly influenced by membrane composition [10, 25–27]. Membrane disruption without pore formation has also been attributed to curvature stresses generated on the extracellular leaflet by AMP binding, with transient transport channels arising through lipid flip-flop events [23, 24]. CL can directly influence these membrane-disruptive processes. In an effort to elucidate the role of CL on pore formation and AMP activity, MD simulations have shown a higher free energy for pore formation with increasing CL content in PG membranes [27]. In another study, CL was found to reduce mechanical deformations induced by aurein 1.2, a 13-residue peptide, due to its intrinsic negative-curvature-inducing property [28]. Despite these insights into downstream membrane disruption, one of the challenges lies in capturing the initial transition of an AMP from an unfolded or partially folded state in solution to a membrane-inserted folded state and determining how membrane composition influences the energetics of this transition.

In this work, we use all-atom molecular dynamics simulations to study the free energy associated with CM15 interactions with the bacterial inner membrane and, in particular, decipher the role played by CL in membrane binding and folding. The 15 residues AMP ‘CM15’ (KWKLFKKIGAVLKVL) is a combination of cecropin, A (1–7 residues) and melittin (2–9 residues). It is a synthetically developed cationic intrinsically disordered peptide with a higher antimicrobial potency and minimal hemolytic activity [29–31]. A study using site-directed spin labelling [32] reported that the peptide is located below the membrane headgroups at an average depth of ∼5Å bound parallel to the membrane plane. Leakage and membrane disruption by CM15 is generally attributed to a toroidal pore-forming mechanism due to its small size relative to the membrane thickness, although no direct evidence has been reported [33]. Few MD simulations have been carried out on CM15 to understand its mechanism of action. In the study by Bennet et al.,[20] the unfolded state of a single peptide in water was found to fold and remain near the lipid headgroup regions of a POPE:POPG bilayer. In an earlier all-atom MD simulation [34] of CM15 in POPC and POPC-POPG membranes, peptide interactions with the anionic POPG containing membranes were stronger, and the peptide had a greater propensity to fold in the POPC membranes. The authors concluded that the lowered electrostatic interactions in the POPC membranes would lower the free-energy barrier for folding. In a distinct departure from studies with phospholipid bilayers, all-atom MD simulations were used to evaluate the free energy of translocation of CM15 through the outer LPS layer of Gram-negative bacteria [35] using only the membrane normal distance as the collective variable. The largest barrier was observed in the highly charged core saccharide region of the lipopolysaccharide membrane. Despite the insights gained from these biased MD simulations, the free energy associated with CM15 membrane binding and folding is yet to be reported.

Describing this transition requires considering two major factors: the relevant degrees of freedom needed to capture both peptide translocation in the membrane and secondary-structure changes, and a realistic description of the bacterial membrane composition [10]. We apply the finite-temperature string method [36, 37] to obtain the free-energy landscape associated with CM15 insertion from an unfolded state in solution to a membrane-inserted folded state. We use a two-dimensional collective-variable space considering both the position of the peptide and *α*-RMSD [38], which has been shown to reliably capture helical structural changes in peptides [38, 39]. To our knowledge, this is the first study to incorporate the secondary structure as a collective variable for AMP-membrane interactions. Previous free energy studies with AMPs interacting with lipid membranes have used, in addition to the distance of the peptide from the membrane, the tilt angle of the peptide with the membrane normal, without explicitly accounting for the secondary structure changes [18, 19, 40].

Our all-atom finite temperature string method overcomes this limitation, allowing us to capture the ubiquitous folding transition associated with AMP membrane binding and insertion. Free energy computations are carried out for membranes with PE:PG (79:21) and PE:PG:CL (75:20:5) to understand the putative role played by CL in influencing the barrier for folding in the membrane. The underlying mechanism follows three key steps: membrane binding of the unfolded peptide, membrane insertion of the unfolded peptide, and folding of the peptide under the lipid headgroup to a phospholipid headgroup parallel state. We observe that the presence of CL prevents the peptide from attaining an *α*-helical state, giving rise to barriers ranging from 2.5 - 8 *k*_*B*_*T* for folding. In the absence of CL, the peptide is found to freely insert below the phospholipid headgroups and spontaneously fold, adopting a membrane parallel state. Small unilamellar vesicle leakage experiments are carried out with various peptide:CL ratios. These experiments illustrate a decrease in leakage with increasing CL content for low peptide concentrations, which we attribute to strong pinning of CM15 with CL preventing downstream lytic events driven by peptide and lipid reorganization. Together, these results illustrate how membrane composition can reshape the free-energy landscape of CM15 insertion and folding, thereby modulating downstream membrane-disruptive activity.

## MATERIALS AND METHODS

The initial structure of the peptide CM15, reported as a micelle-bound *α*-helical structure (Figure S1), with amino acid sequence ‘KWKLFKKIGAVLKVL’ was taken from the RCBS PDB database (2JMY) [41]. CM15 is made up of mainly hydrophobic residues with one third being positively charged lysine residues (Figure S1). We performed NPT simulation of the peptide in an aqueous environment with 3561 TIP3P water molecules [42], and 0.15 M NaCl salt concentration. The constant pressure (1 bar-isotropic) and temperature (303.15 K) were maintained using Parrinello-Rahman barostat and Nosé-Hoover thermostat. Bacterial inner membrane consisting of 75% 1,2-dioleoyl-sn-glycero-3-phosphoethanolamine (DOPE), 20% 1,2-dioleoyl-sn-glycero-3-[phospho-rac-(1-glycerol) (DOPG), and 5% tetraoleoyl cardiolipin (TOCL1) was prepared using CHARMM-GUI server [43–45]. Similarly, a membrane with 79% DOPE and 21% DOPG was also prepared using CHARMM-GUI server. For the MD simulations, different initial states, such as the folded and unfolded peptide states, exposed to the membrane-water interface and placed below the membrane headgroups, were prepared for both the membranes. Periodic boundary conditions were applied in all three directions. Simulations were performed using a leapfrog integrator with an integration time step of 2 fs. The Nosé-Hoover thermostat [46] (coupling constant of 1.0 ps) and Parrinello-Rahman barostat [47] (semi-isotropic pressure coupling with a coupling constant of 5.0 ps) were used to perform constant temperature (303.15 K) and constant pressure (1 bar) simulations. The isothermal compressibilities for the barostat were set to *k* _*xy*_ = *k* _*z*_ = 4.5 10 ^− 5^ bar ^− 1^. Linear constraint solver (LINCS) [48] was used for constraining hydrogen bonds. The Particle Mesh Ewald (PME) algorithm [49] was used to compute electrostatic interactions with a 1.2 nm cutoff. The van der Waals interactions were smoothly switched to zero between 1.0 and 1.2 nm. A complete list of the simulations performed in the study is given in Table S1. All simulations were performed using GROMACS version 2022.5 [50] patched with PLUMED version 2.9 [51, 52].

### Finite temperature string method simulation details

We used two collective variables, the *z* coordinate of the distance between the center-of-mass of the peptide and the center-of-mass of the bilayer (denoted as *d*_*z*_) to account for the position of the peptide with respect to the membrane, and the *α*-RMSD to account for the helical content of the peptide (see Figure S2). The *α*-RMSD is defined from the co-ordinates of six consecutive residues of the peptide during the simulation compared with the co-ordinates of the ideal helix [38]. The *α*-RMSD is defined as,

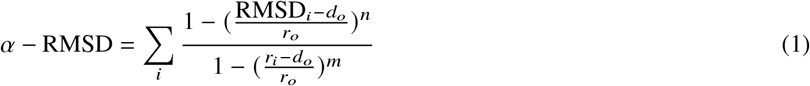

where RMSD_*i*_ is the RMSD of every six consecutive residues, *d*_*O*_ and *r*_*O*_ control the smoothness of the function and are considered as 0 and 0.08 in our study, respectively. Values of the power *n* and *m* are considered as 8 and 12, respectively, same as in the original work [38]. The reference structure for the RMSD computations is a helical crystal structure or a self-prepared reference structure. In our case, we consider the helical crystal structure as the reference state, which has an unstructured end terminus (Figure S1).

We utilize the finite temperature string (FTS) method [36] to capture the CM15 transition and perform the weighted histogram analysis method (WHAM) [53] to compute the free energies. In the biased simulation used in the string method, a force constant of 500 kJ mol^−1^ was applied to both collective variables. During the string evolution, 1 ns of simulations were performed at each point in every iteration until the string converged. The convergence of the string was monitored using the error, *E* defined using,

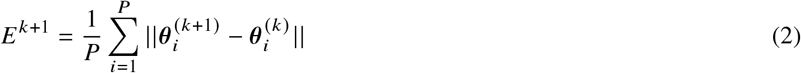

where *E* ^*k* +1^ is the error at iterate, *k, P* is the number of points on the string, with each point denoted by subscript *i*. The *θ* represents the 2D vector of collective variables, *d*_*z*_ and *α*-RMSD. The error, *E* ^*k* +1^ for the PE:PG membrane for the string iterations is plotted in (Figure S3A). The initially constructed 11-point string converged in 16 iterations. The number of points was then increased to 21 after the 22^*nd*^ iteration to refine the string, and convergence was achieved with an additional 16 iterations. For the PE:PG:CL, we obtained two string evolution paths, which we refer to as S-I and S-II. For the PE:PG:CL membrane (Figure S3B), convergence for the S-I string was obtained after the 15th iteration. The S-II string is derived by using a different initial configuration for the extracellular endpoint, and retracing the S-I path to generate initial configurations for the string evolution. Convergence is observed after 9 (Figure S3C) iterations. An additional 200 ns of umbrella sampling simulations were performed at each point of the converged paths for the PE:PG membrane and the S-II evolution, and 180 ns of umbrella sampling on each point for the S-I case. All simulations were carried out at a temperature of 310.15 K to obtain a converged free-energy landscape along the transition path. The string method umbrella sampling computations resulted in a cumulative 12.4 *μ*s of all-atom simulation time.

The 1D free energy profile was reported on the points on the final reported string, calculated from the membrane-exposed to membrane-inserted folded state. Additional convergence checks were performed on the free energy evolution. In Figure S4, the free energy for all three strings was found to be well-converged by 180 ns of umbrella sampling simulations at each point. The bootstrapped error analysis was carried out using the block bootstrapping method. The trajectory obtained from each umbrella sampling window was split into 50 blocks. A new data set was then constructed by random resampling of the blocks with replacement for each window. Using this new dataset, WHAM calculations were performed as described above. This process was then repeated for 500 resampling cycles. To evaluate the error from these resampled free energies, the standard deviation and the average energy value were calculated across the different samples.

### Simulation analysis

Gromacs in-built commands were used to compute root mean square deviation (RMSD) and radius of gyration. The MDAnalysis Python package was used for post-processing of the trajectories. Time trajectories of secondary structure changes were monitored using the VMD timeline features with the STRIDE software. For the lipid molecules occupancy analysis on the residue of the peptide, contact between two atoms was assumed to occur if the distance was less than a cutoff distance of 0.5 nm [54, 55]. In the residue-wise occupancy analysis of the lipid molecules, a contact was counted if any atom of the lipid came within a 0.5 nm cutoff distance of any atom of the peptide residue [54, 55]. Thus, a high number of contacts would indicate a strong interaction that would not be captured by the center-of-mass distance between the residue and the lipid/sterol criteria. The lipid contact for residue, *r* is defined using,

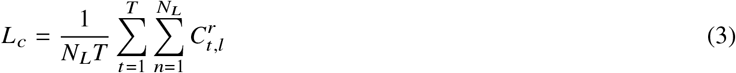

where *t* is the time index, *T* is the total simulation time, *n* is the index for number of lipid molecules, *N*_*L*_ is the total number of lipid molecules. 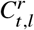 denotes the total number of contacts formed between *r*^*th*^ residue and *l*^*th*^ lipid molecule at time *t* and is evaluated using,

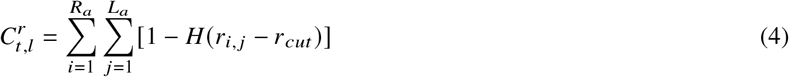

where *H* (*x* − *x*_o_) is the Heaviside step function whose value is 0 when *x < x*_*o*_ and 1 when *x ≥ x*_*o*_. *R*_*a*_ is the number of atoms in a residue and *L*_*a*_ is the number of lipid atoms. Final occupancy values were also normalized with the number of tails of a lipid molecule.

The hydration shell per number of total molecules analysis for water molecules and a similar analysis for lipid molecules, called lipid occupancy per number of molecules, is done using ‘around’ feature of the MDAnalysis Python package [56, 57]. This provides the number of molecules in a specified cutoff away from the peptide, where the cutoff boundary is constructed using the surface accessible surface area (SASA). We used a cutoff of 0.35 nm. The curvature analysis was performed using the ‘MembraneCurvature’ Python package based on the method proposed by Yesylevssky et al. [58], which uses a grid-based method to construct the height-field function.

## EXPERIMENTAL METHODS

### Liposome Preparations

25 mg/ml stock solutions of the lipid components 1,2-di-(9*Z*-octadecenoyl)-*sn*-glycero-3-phosphoethanolamine (DOPE), 1,2-di-(9*Z*-octadecenoyl)-*sn*-glycero-3-phospho-(1 ′ -*rac*-glycerol) (DOPG), and 1,1 ′,2,2 ^′^ -tetra-(9*Z*-octadecenoyl) cardiolipin (CL) (Avanti Polar Lipids, USA) were dissolved in chloroform and stored at −20 ° C in clean glass vials. The lipid mixtures were added proportionally in clean glass vials to give a final concentration of 3 mM upon hydration. The ratio of DOPG:DOPE was kept constant at 75:20 in all vesicle formulations, and the CL percentage was varied from 0% to 20% (Table S2). The lipid mixture was dried using a gentle nitrogen stream to form a lipid film on the glass wall surface. The dried lipid film was kept under vacuum for 1 hour to remove any traces of chloroform. The lipid film was hydrated using 50 mM Sulforhodamine B (SRB) (Merck) dissolved in phosphate-buffered saline (PBS, pH 7.4) at 37 ° C for two hours and then vortexed vigorously to obtain multilamellar vesicles. The vesicles were freeze–thawed using liquid nitrogen for five cycles and extruded using 0.1 *μ*m membranes to obtain small unilamellar vesicles (SUVs). The SUVs were purified using gel filtration (Sephadex G-25). The vesicle fractions were pooled together and stored at 4 °C. The vesicle size was measured using dynamic light scattering (NanoBrook ZetaPALS, Brookhaven Instruments), yielding diameters of 136.6±5.6 nm (Figure S5).

### Vesicle leakage assay

The SRB fluorescence was measured using 559 nm excitation and 577 nm emission with a 5 nm slit width. Vesicles were diluted 100-fold in PBS, and the total dye content was measured by adding Triton-X (final concentration 1%) to the solution. The vesicle volumes were normalized to ensure equal SRB fluorescence rise for all lipid compositions using spectrophotometer (Agilent Cary Eclipse). Vesicle leakage upon addition of CM15 (NovoPro Biosciences, China) at different peptide-to-lipid ratios (Table S3) was measured for 20 minutes in triplicate in a 96-well black fluorescence plate (Bio-Rad Laboratories, USA) using a Thermo Fisher Varioskan microplate reader. The leakage kinetic data were normalized to the total Triton-X fluorescence to obtain the fractional leakage:

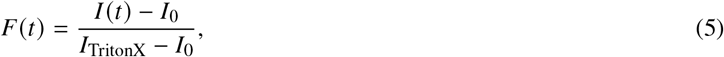

where *F(t)* is the fractional leakage at time *t, I(t)* is the SRB intensity at time *t, I*_*0*_ is the intensity of untreated vesicles, and *I*_TritonX_ is the intensity of SRB after Triton-X treatment. The normalized fractional leakage data are given in Supplementary Figure S6. Due to inherent instrument delay times (∼ 10 s) the short time leakage data is not captured. The *t*_1/2_ values were calculated for peptide-to-lipid (P:L) ratios of 1:20 directly from each raw normalized leakage trace (Figure S6) by identifying the time point at which the signal reached half of its maximum value. We were unable to reliably extract similar half-time leakage values for higher P:L ratios.

## RESULTS

### CM15 unfolds in an aqueous environment

The crystal structure of CM15 was reported as an *α*-helix in the micelle-bound membrane mimicking environment [41]. Restraint free MD simulations of solvated CM15 in 0.15 M NaCl in the NPT ensemble show that the initially folded state rapidly unfolds in the bulk aqueous environment. Two independent simulations (1 *μ*s each) illustrate that the peptide rapidly unfolds within the first 100 ns and remains unfolded over the rest of the simulation (Figure S7A). The unfolding of CM15 in an aqueous environment is consistent with previously published reports [35, 41]. The secondary structure analysis validates that the *α*-RMSD is a reliable metric to quantify the helical content in the peptide (Figure S7B).

### Cardiolipin prevents spontaneous folding of CM15 upon membrane binding in unrestrained MD simulations

To understand the membrane insertion and folding mechanisms, we initially carried out unbiased MD simulations with an unfolded CM15 molecule placed in the proximity of the PE:PG lipid headgroups. The results are illustrated for a 4 *μ*s simulation in Figure 1. Snapshots illustrate the initial unfolded peptide in the proximity of the membrane with subsequent folding upon membrane entry (Figure 1A). The time evolution of the *α*-RMSD and d_*z*_ measured from the bilayer mid-plane to the average headgroup *z*-plane is illustrated in Figure 1B. CM15 inserts into the membrane in an unfolded state and folds into a predominantly helical configuration at around 1 *μ*s, remaining mostly folded over the next 3 *μ*s with *α*-RMSD > ∼ 1.5. Over this period, the peptide remains below the phospholipid headgroups with a mean value of d_*z*_ = 1.807 ± 0.182 nm (averaged over the last 3 *μ*s). The 2D histogram map (Figure 1C) of the *α*-RMSD versus d_*z*_ for the final 2 *μ*s of the 4 *μ*s simulation illustrates the stability of the folded states of the peptide upon membrane entry. We also evaluated other secondary structure motifs, which confirm that the peptide remained predominantly in the folded state (Figure S8A). Similar trends were observed for an independent 2 *μ*s simulation (see Figure S9), where we observe folding upon entry into the membrane. Similar trends were observed in MD simulations by Bennet et al., with a single CM15 molecule placed on the leaflet of a PE:PG membrane [20]. We next contrast these trends with the PE:PG:CL membrane.

**Figure 1:**
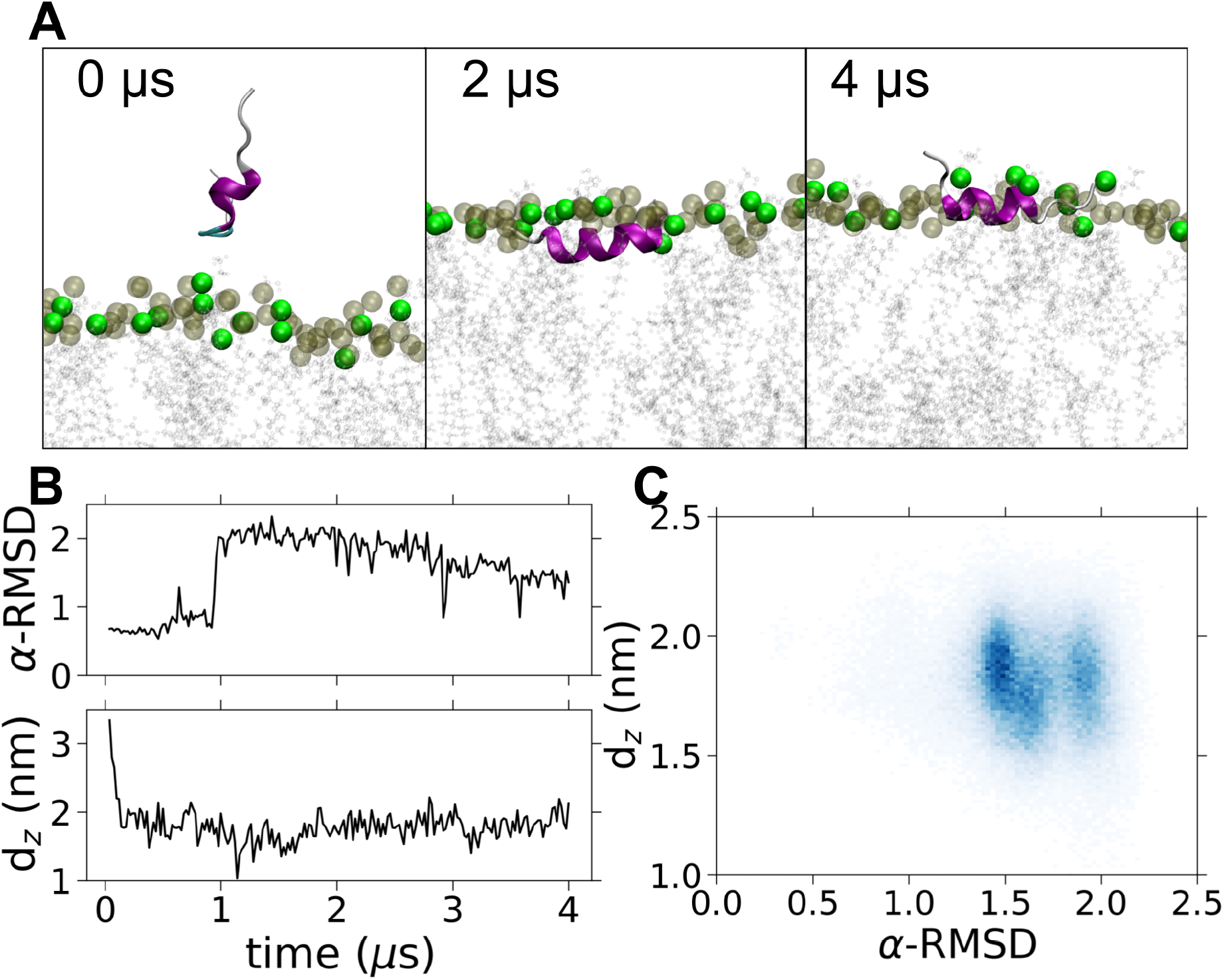
Unrestrained simulations of CM15 in a PE:PG membrane. (A) Snapshots illustrating the orientation and secondary structure adopted by CM15 at different stages of a 4 *μ*s all-atom MD simulation. (B) Time evolution of the *α*-RMSD and d_*z*_ collective variables. (C) 2D histogram of d_*z*_ versus *α*-RMSD depicting the different configurations, sampled over the final 2 *μ*s of simulation. CM15 inserts in an unfolded state, adopting a folded configuration beneath the lipid headgroups beyond 1 *μ*s. Green beads-PG lipid headgroups; gray beads-PE lipid headgroups. Lipid tails and water are not shown for clarity.

In the PE:PG:CL membrane (Figure 2), CM15 sampled predominantly unfolded states with a low *α*-RMSD *<* 0.5 (Figure 2B). The peptide was also found to be located predominantly below the phospholipid head groups with a mean d_*z*_ value of ∼ 1.9 nm (Figure 2B). The 2D histogram of the *α*-RMSD and d_*z*_ for the final 2 *μ*s of the 4 *μ*s simulations is illustrated in Figure 2C. Similar trends were observed for an independent simulation of 2*μ*s (Figure S10). These MD simulations illustrate that although CM15 is able to spontaneously insert into the membrane in an unfolded state and remain colocalized with the membrane headgroups, there exists a distinct barrier to folding in the PE:PG:CL membrane (Figure S8B). This is in sharp contrast to the PE:PG membrane, where CM15 spontaneously folded upon membrane insertion. Although CL is only present in 5 mol % of the membrane composition, our simulations suggest a dominant role in preventing CM15 from spontaneously accessing a folded state upon membrane binding. In order to assess the extent of the free energy barriers and quantify the energetics associated with CL, we carry out string method free energy simulations in both membranes.

**Figure 2:**
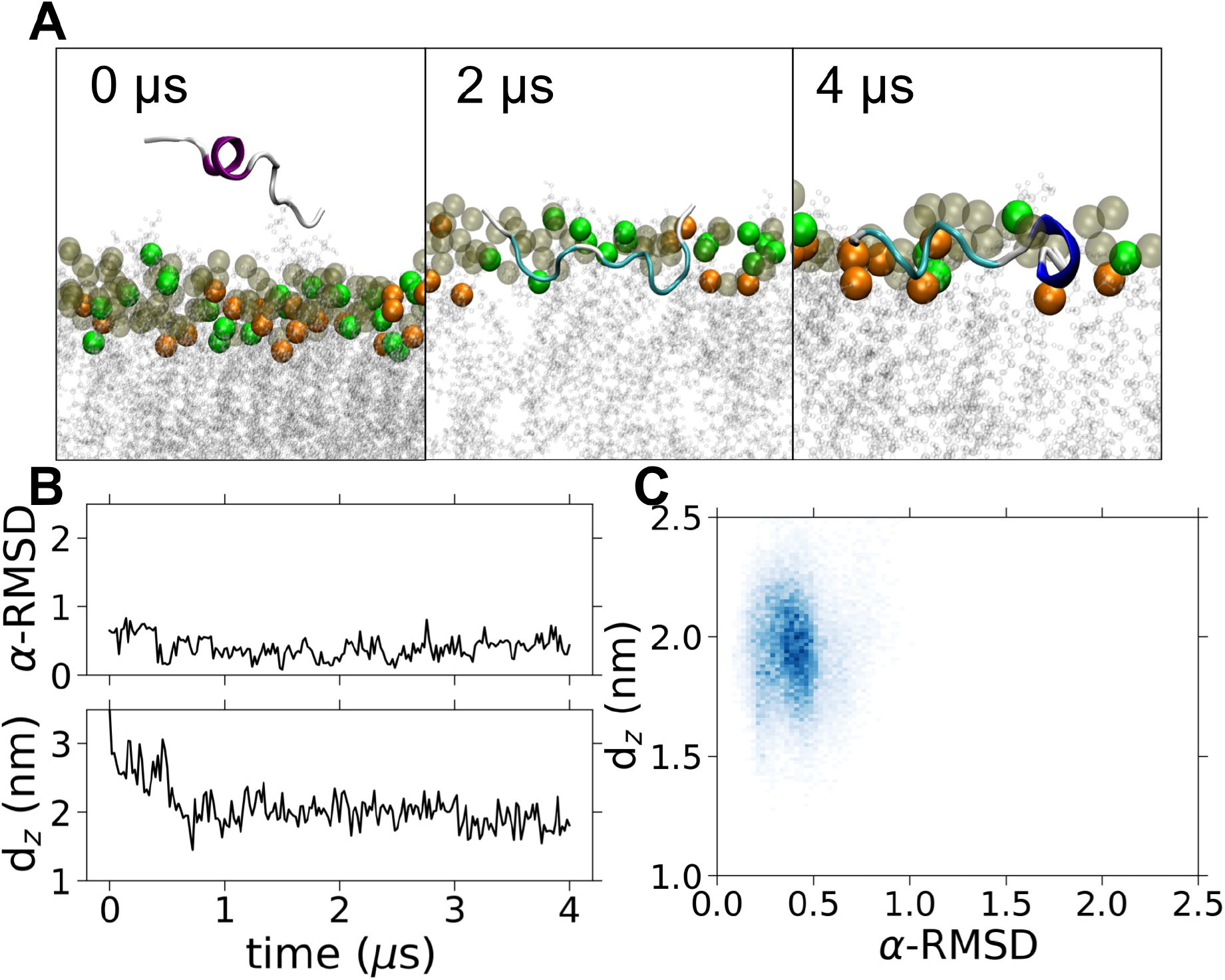
Unrestrained simulations of CM15 in a PE:PG:CL membrane. (A) Snapshots illustrating the orientation and secondary structure of CM15 at different stages of a 4 *μ*s all-atom MD simulation. (B) The time evolution of *α*-RMSD and d_*z*_. (C) 2D histogram of d_*z*_ versus *α*-RMSD, sampled across the final 2 *μ*s. CM15 inserts into the membrane in an unfolded state and remains unfolded over the course of the simulation. Green beads-PG lipid headgroups; gray beads-PE lipid headgroups, orange beads-CL lipids. Lipid tails and water are not shown for clarity.

### Free-energy landscape reveals spontaneous membrane insertion and folding of CM15 in the PE:PG membrane

Based on the MD simulations discussed above, we identify the two endpoints for the string method free-energy simulations. For the PE:PG membrane, the two end points are, respectively, an unfolded state in the extracellular volume with *α*-RMSD = 0.1 and d_*z*_ = 3.0 nm, and a folded membrane-parallel state with *α*-RMSD = 2.1 and d_*z*_ = 1.7 nm.

The Figure 3 illustrates the results from the string method free energy computations for the PE:PG membrane. The converged path in the d_*z*_-*α*-RMSD collective variable space is illustrated in Figure 3A and the corresponding free energy variation along the path in Figure 3B. Selected snapshots along the string illustrate the different conformation states sampled by the peptide (Figure 3C). For the PE:PG membrane, the free energy continuously decreases along the path as CM15 enters the membrane from the extracellular environment. The d_*z*_ decreases from 3 nm to 2.75 nm, bringing the peptide in contact with the phospholipid headgroups of the membrane (point 2 in Figure 3A,B). At this point the peptide is predominantly in an unfolded state, and membrane binding of the positively charged peptide with the negatively charged bacterial membrane is accompanied with a large free energy decrease of ∼ 7*k*_*B*_*T* . We observe a steady decrease in the free energy along the path as the peptide inserts deeper into the headgroup region of the membrane, while the *α*-RMSD shows very little change during this insertion stage (first 8 points along the path). Beyond *d*_*z*_ = 2.1 nm, further insertion is accompanied by an increase in *α*-RMSD, eventually reaching a minimum at *α*-RMSD = 1.1 with *d*_*z*_ = 1.85 (point 13 on the string). To transition to a fully folded state, a small free energy barrier of ∼ 3 *k*_*B*_*T* is observed. This suggests there is a free energy basin around the minima where the peptide can access different conformations with *α*-RMSD varying from ∼0.6 to ∼ 2.1 without encountering any significant barriers. This supports the behavior observed in unbiased MD simulations, in which the peptide remains folded with only a small variation in *d*_*z*_. Selected snapshots along the string illustrate the various stages along the path, and we identify the following states as illustrated in Figure 3C. A surface bound unfolded state, *S*_*u*_, a membrane bound unfolded state, *M*_*bu*_ a membrane inserted unfolded state, *M*_*iu*_, and a membrane inserted folded state, *M*_*if*_ . The *α*-RMSD varies from 1 to 2.1, distinguishing the *M*_*iu*_ state from the fully folded *M*_*if*_ state.

**Figure 3:**
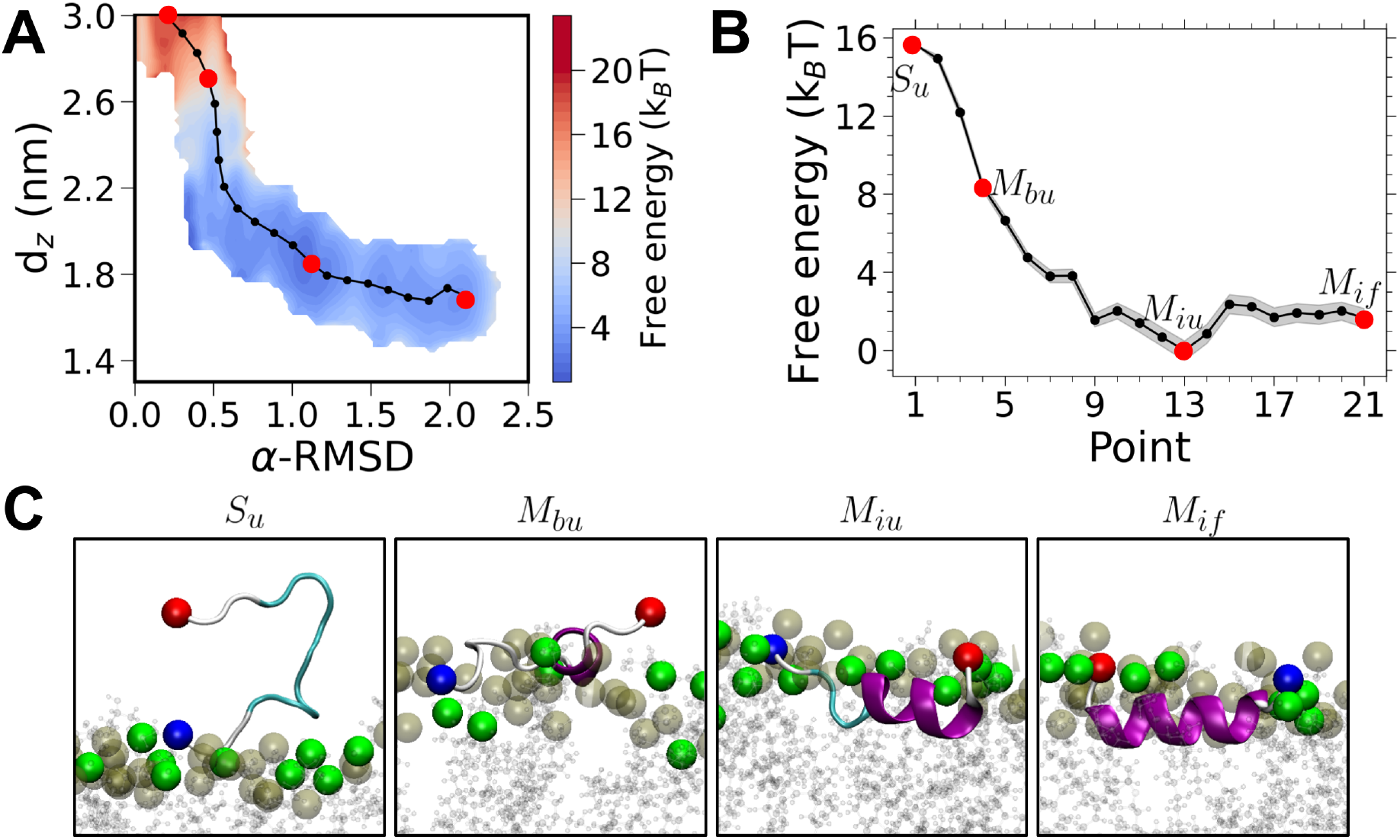
Free energy for CM15 insertion pathway in the PE:PG membrane. (A) 2D free energy along the converged string (black) for the PE:PG membrane and (B) corresponding 1D free energy profile along the path with different states marked in red dots, S_*u*_ - solvent unfolded, M_*bu*_ - membrane bound unfolded, M_*iu*_ - membrane inserted unfolded and M_*if*_ - membrane inserted folded (see Figure 6). (C) Snapshots illustrating states at points marked in red along the string. Color scheme for snapshots (C), red sphere (N-terminus) and blue sphere (C-terminus); green beads-PG lipid headgroups; gray beads-PE lipid headgroups. Lipid tail and water molecules are not shown for clarity.

### Cardiolipin introduces distinct free-energy pathways for CM15 insertion and folding

In order to study the effect of cardiolipin, we compute the free energy of CM15 insertion and folding using the string method. We select the two endpoints for the PE:PG:CL membrane from the previously discussed unbiased simulation: a partially folded state with *α*-RMSD = 0.8 and d_*z*_ = 2.5 nm, and a folded membrane-parallel state with *α*-RMSD = 2.1 nm and d_*z*_ = 1.8 nm. The corresponding results of the free-energy computations for the PE:PG:CL membrane are shown in Figures 4 and 5, where we illustrate two distinct paths that emerge during the string evolution. This is indicative of the rugged nature of the free energy landscape encountered by the peptide in the presence of CL. We denote these two different string evolution paths as S-I and S-II. In S-I (Figure 4), the initial *S*_*u*_ state has an *α*-RMSD of 0.8. The peptide present on the membrane surface fully unfolds upon insertion into the membrane (*α*-RMSD from 0.8 to 0.08) to give rise to the *M*_*bu*_ state as illustrated in Figure 4C with the free energy landscape having a broad minima around this low value of *α*-RMSD (Figure 4B). Beyond this point (point 6) on the path, we observe a large barrier (∼ 10 *k*_*B*_*T* ) for further insertion and initiation of peptide folding. In this phase the *α*-RMSD varies from 0.1 to 0.5 and d_*z*_ from 1.8 to 1.5 nm (point 6 to 10 in Figure 4A) where the *M*_*iu*_ state is formed. The free energy for the subsequent folding of the inserted peptide to *α*-RMSD = 2.0, is downhill. In contrast to the peptide binding and folding in the PE:PG membrane (Figure 3A,B), where insertion and folding occur contiguously, the presence of CL results in a barrier for membrane insertion and positioning of the peptide below the headgroups of the membrane. Folding is favored once this positioning along the d_*z*_ co-ordinate is complete. The snapshots shown in Figure 4C elucidate this insertion and subsequent folding behavior of the peptide along the string.

**Figure 4:**
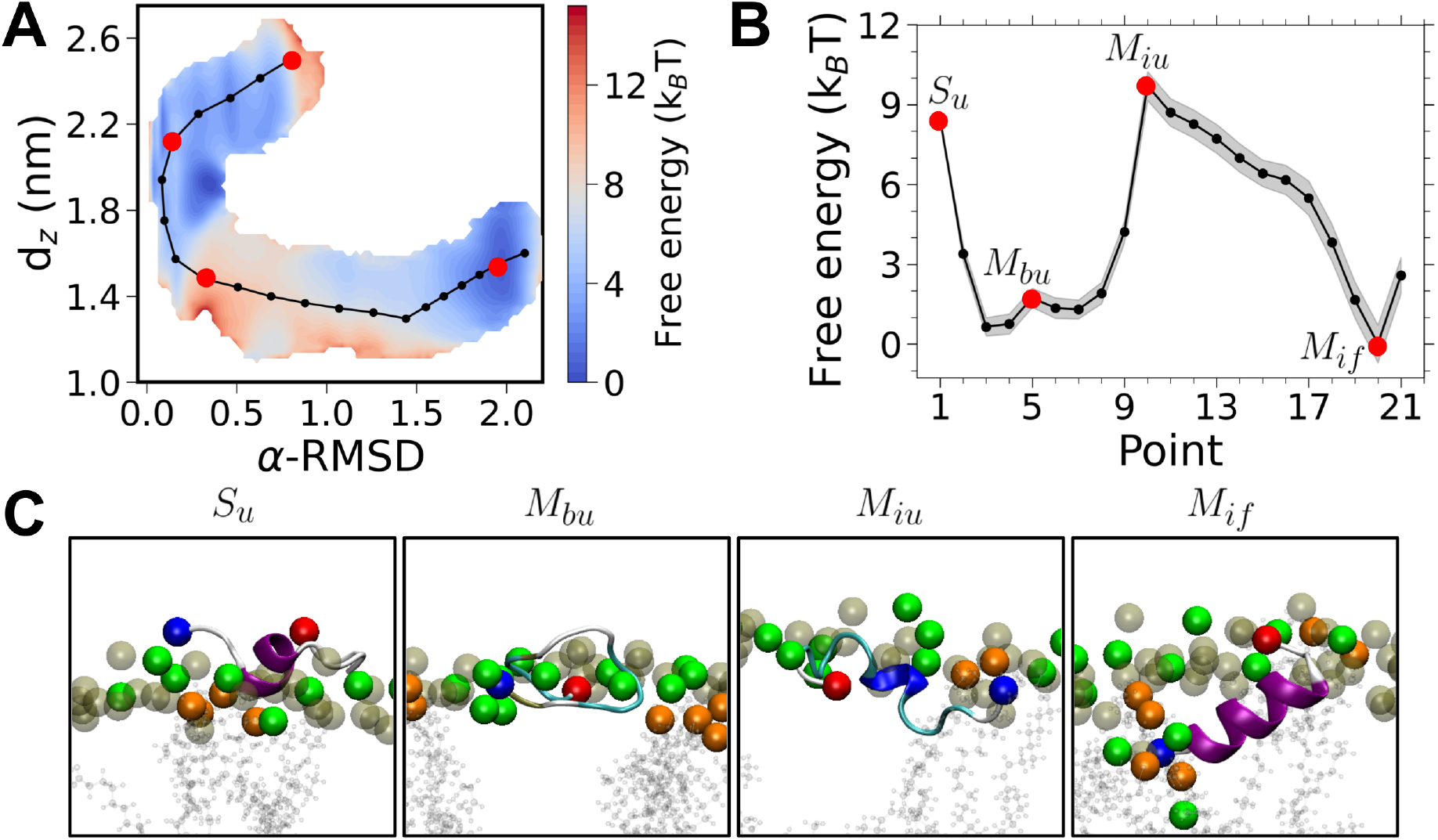
Free energy for CM15 insertion pathway S-I in the PE:PG:CL membrane. (A) 2D free energy contour plot (black). (B) Corresponding 1D free energy profile along the path with different states sampled at the points marked red along the string (see snapshots in C), S_*u*_ - solvent unfolded, M_*bu*_ - membrane bound unfolded, M_*iu*_ - membrane inserted unfoled and M_*if*_ - membrane inserted folded (see Figure 6). Color scheme for snapshots (C), Red sphere (N-terminus) and blue sphere (C-terminus); green beads, PG lipid headgroups; gray beads, PE lipid headgroups, orange beads, CL lipids. Lipid tail and water molecules are not shown for clarity.

**Figure 5:**
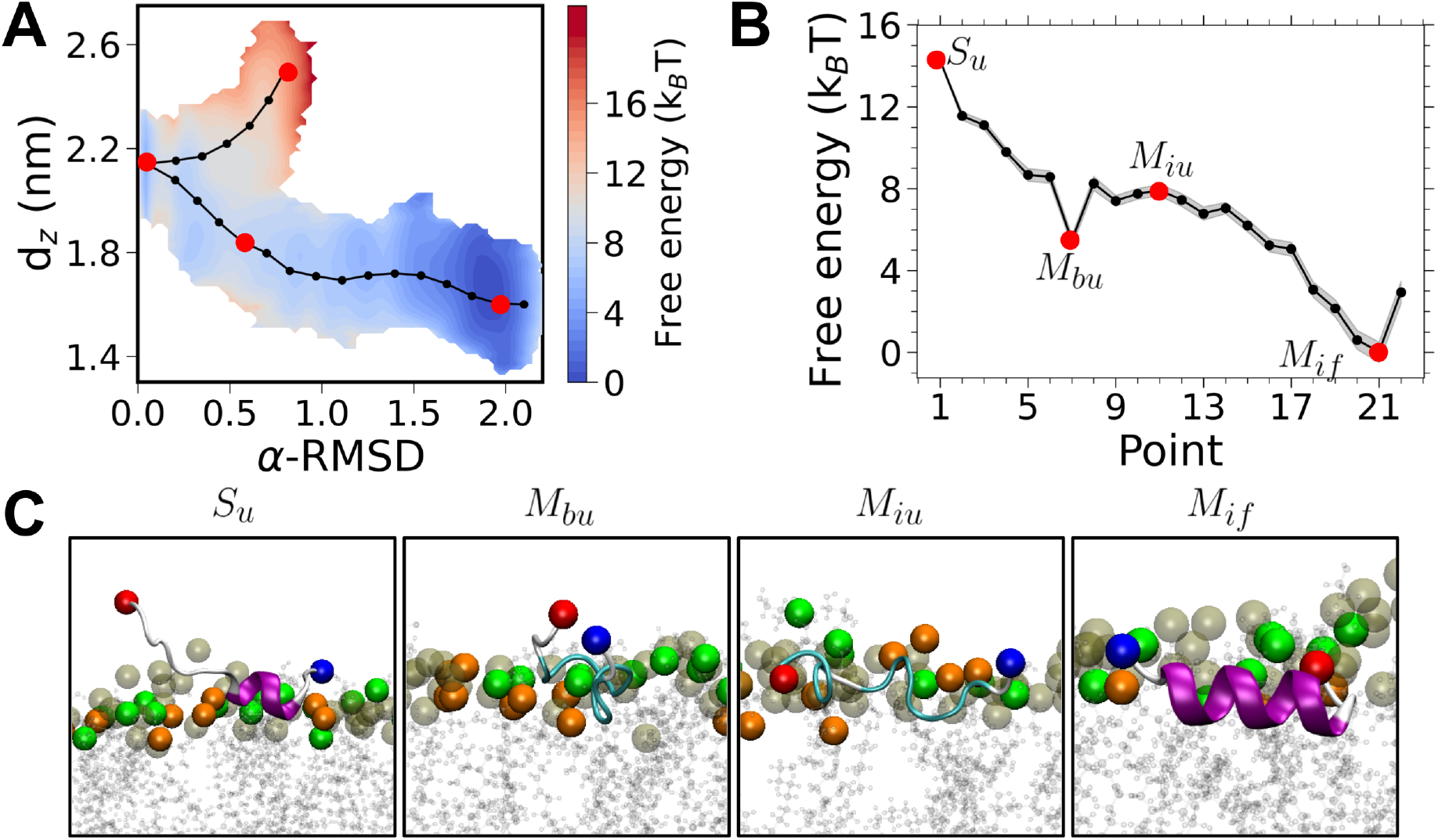
Free energy for CM15 insertion pathway S-II in the PE:PG:CL membrane. (A) 2D free energy along the converged string (black). (B) Corresponding 1D free energy profile along the path with different states sampled at the points marked red along the string (see snapshots in C), S_*u*_ - solvent unfolded, M_*bu*_ - membrane bound unfolded, M_*iu*_ - membrane inserted unfoled and M_*if*_ - membrane inserted folded (see Figure 6). In contrast to the S-I pathway (Figure 4), a small barrier to transform from an unfolded state (M_*bu*_) to the membrane inserted folded state (M_*if*_ ) is observed. Color scheme in snapshots (C); red sphere (N-terminus) and blue sphere (C-terminus); green beads, PG lipid headgroups; gray beads, PE lipid headgroups, orange beads, CL lipids. Lipid tail and water molecules are not shown for clarity.

In S-II (Figure 5), the partially folded peptide (*S*_*u*_) present on the membrane surface completely unfolds to insert into the headgroup region of the membrane (*α*-RMSD going from 0.8 to 0.08) to form the *M*_*bu*_ state. This transformation is similar to what was observed in S-I (Figure 4). Differences between the pathways emerge beyond this unfolded state. In S-II, we observe a free energy minima corresponding to the *M*_*bu*_ state as observed in S-I. The free energy decrease from the *S*_*u*_ to the *M*_*bu*_ state is similar for both S-I and S-II, with a decrease of ∼ 7 *k*_*B*_*T* . However, in contrast to the S-I landscape, the *M*_*bu*_ state for S-II corresponds to a weak local minima (Figure 5B). Subsequently, the peptide undergoes membrane insertion from the *M*_*bu*_ state to the partially folded *M*_*iu*_ state. During this membrane insertion state, *d*_*z*_ decreases to ∼ 1.8 nm with an increase in the *α*-RMSD value of 0.6. A weak local barrier of about 2 *k*_*B*_*T* is observed during this transition. Once the peptide has reached the *M*_*iu*_ state, it is able to spontaneously fold, and the *α*-RMSD increases to ∼ 2 with a further modest insertion into the membrane. Subsequent folding of the peptide occurs spontaneously, and the free energy minimum occurs at *α*-RMSD 2.0 and *d*_*z*_ of ∼ 1.6 nm, similar to the final state obtained in S-I (Figure 4). The presence of barriers for folding in the presence of CL is consistent with our unbiased MD simulations, where we did not observe spontaneous peptide folding events across multiple microsecond-long simulations (Table S1).

Comparing S-I and S-II strings, we observe differing free energy landscapes that emerge along the path. S-I shows a large free energy barrier for the initial folding of the peptide in the interior of the membrane from an unfolded state. However, this is not observed in S-II, where the initial folding transition occurs closer to the membrane surface, and a broad plateau region then follows in the free energy landscape. We note that the presence of the barrier in the S-I pathway is associated with the deeper insertion of the peptide into the membrane (*d*_*z*_ ∼ 1.5 nm, Figure 4A,B) suggesting that unfolding of the peptide is facilitated when it is in closer proximity to the phospholipid headgroups as seen in the S-II pathway (*d*_*z*_ = 1.8 nm, Figure 5A,B). Beyond the *M*_*iu*_ state, folding is downhill in both cases, and the respective overall free energy minima occur at the *α*-RMSD of 2.0 and *d*_*z*_ of ∼ 1.6 nm. Despite the absence of large barriers, the free energy variation along the transition is inherently rugged. The free-energy pathways in the PE:PG:CL membrane clearly indicate the presence of a modified landscape with barriers induced for CM15 folding in the presence of CL. Although CL has been shown to increase the energetic penalty for the formation of aqueous pores in PG:CL membranes [27], our study illustrates for the first time a direct influence of CL on the free energy for a peptide folding pathway in bacterial membranes. The challenge lies in providing a molecular interpretation for the observed changes in CM15 folding induced by CL. After a brief summary of the free energy changes for the two membranes, we explore several factors that provide insights into the putative role of CL in CM15 binding and conformational transitions.

### Comparison of free-energy landscapes reveals cardiolipin-induced kinetic bottlenecks in CM15 insertion and folding

Based on the free energy analysis, we propose a canonical pathway for CM15 to bind and fold in the membrane as illustrated in Figure 6. The reaction pathway for these states is given by

**Figure 6:**
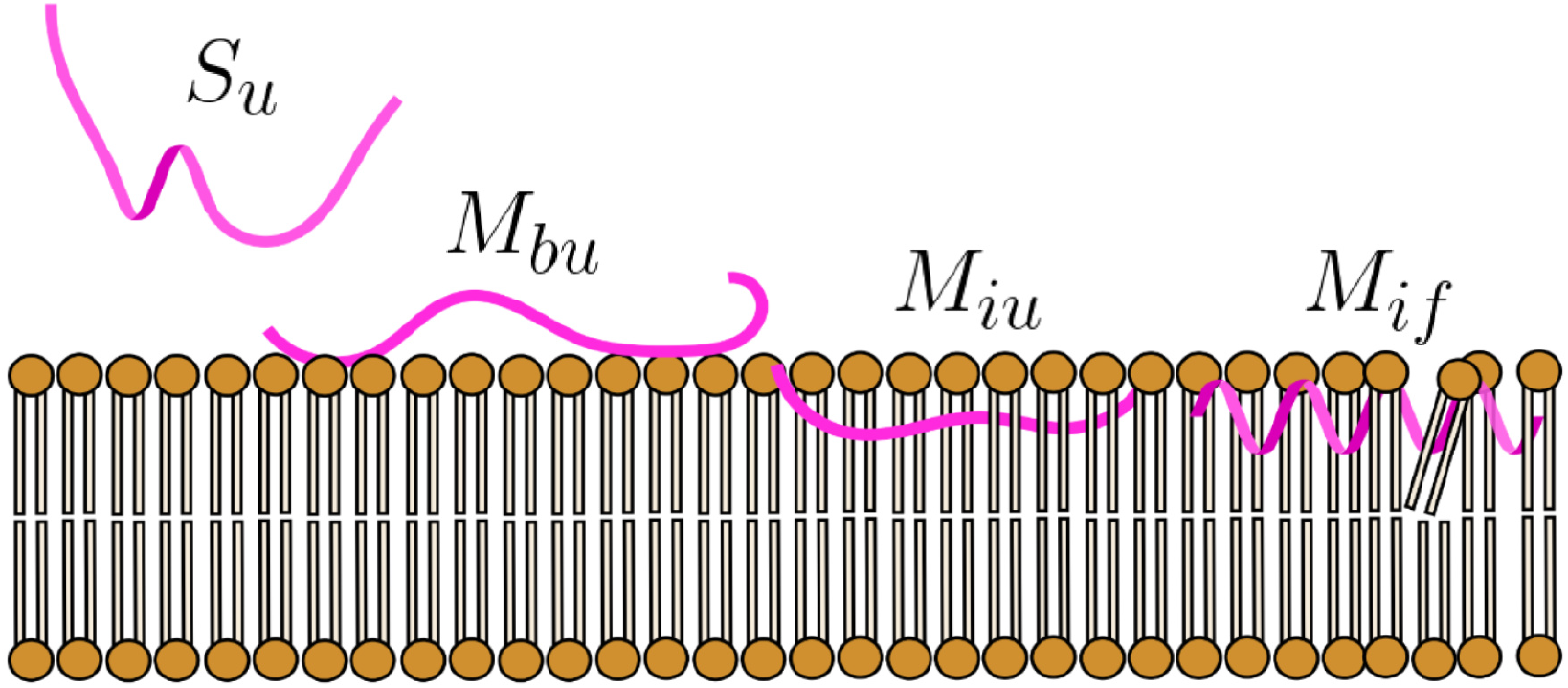
Schematic of various peptide states. S_*u*_-unfolded state in the solvent, M_*bu*_-membrane bound unfolded state, M_*iu*_- membrane inserted unfolded state and M_*if*_ is the membrane inserted folded state.

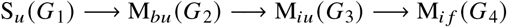

and the free energy associated with each state is denoted as *G*_*i*_,i = 1.. .,4, respectively. The values of the free energy differences for these different states for the two membrane systems are given in Table 1. The transition from S_*u*_ to M_*bu*_ is spontaneous and downhill in the free energy landscape for both membranes with a favorable free energy, Δ*G*_1_ varying between - 6.8 to -8.8 *k*_*B*_*T* .

**Table 1:**
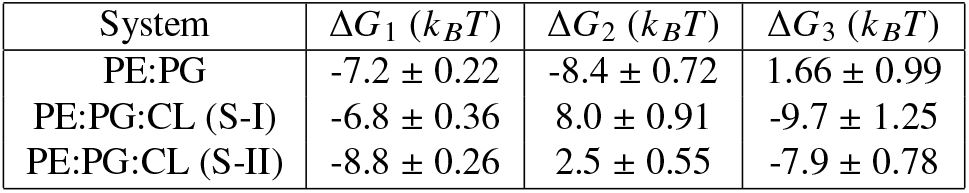
Free energy changes between different states for CM15 membrane entry and folding. Free energy differences Δ*G*_*i*_ are reported between the four different states illustrated in Figure 6. These state are S_*u*_ (*G*_1_), M_*bu*_ (*G*_2_), M_*iu*_ (*G*_3_) and M_*if*_ (*G*_4_). ω*G*_1_ = *G*_2_ − *G*_1_, ω*G*_2_ = *G*_3_ − *G*_2_ and ω*G*_3_ = *G*_4_ − *G*_3_

| System | $\Delta G_1$ ($k_B T$) | $\Delta G_2$ ($k_B T$) | $\Delta G_3$ ($k_B T$) |
| --- | --- | --- | --- |
| PE:PG | $-7.2 \pm 0.22$ | $-8.4 \pm 0.72$ | $1.66 \pm 0.99$ |
| PE:PG:CL (S-I) | $-6.8 \pm 0.36$ | $8.0 \pm 0.91$ | $-9.7 \pm 1.25$ |
| PE:PG:CL (S-II) | $-8.8 \pm 0.26$ | $2.5 \pm 0.55$ | $-7.9 \pm 0.78$ |

The energetics is driven by the binding of the cationic CM15 to the anionic bacterial membrane. For the PE:PG membrane, complete insertion to assume an unfolded state below the headgroups (M_*iu*_) is favorable with an associated free energy change, Δ*G*_2_ = 8.4 *k*_*B*_*T* . In contrast, for the PE:PG:CL membrane, Δ*G*_2_ = 8.0 *k*_*B*_*T* for the S-I pathway and 2.5 *k*_*B*_*T* for the S-II pathway with a weak barrier. The large barrier for S-I is linked to the deeper positioning of the peptide in the lipid milieu. The value of Δ*G*_3_ for complete folding to the M_*if*_ state for the PE:PG membrane is 1.66 *k*_*B*_*T* . The corresponding free energy, Δ*G*_3_, for both the S-I and S-II pathways is favorable, ranging between -7.9 to -9.7 *k*_*B*_*T* . In the absence of CL, the *M*_*iu*_ state is partly folded and a small barrier (1.66 *k*_*B*_*T* ) is observed for complete folding (Figure 3). For the PE:PG:CL membranes, the presence of the free energy barrier for the S-I pathway (Δ*G*_2_ = 8.0) will directly result in a retarded rate for the transition. In contrast, for the S-II pathway, a distinct plateau in the free energy is observed in the vicinity of the *M*_*bu*_ state (Figure 5B), notwithstanding the small change in (Δ*G*_2_ = 2.5), which should be overcome with thermal fluctuations. This free-energy topology with a plateau will also reduce the folding rate, controlled by diffusional exploration in the collective variable phase space in this region. Hence, although the pathways S-I and S-II have distinct topological features, both landscapes will result in a reduction of the rates associated with the overall folding transition [59, 60]. The plateau-like landscapes have also been shown to be associated with large conformational entropy near the fully folded states [61]. In the next section, we compute several metrics to understand the differences in the free energy landscapes.

### *α*-helicity progression sheds light on the difference in free energy of CM15 folding

In this section, we examined the progression of *α*-helicity for each residue of the peptide across all points along the string. This progression is shown in Fig 7, and illustrates the fraction of simulation time spent by each residue of the peptide as an *α*-helix. For the PE:PG membrane (Fig 7A), we see a uniform increase in the *α*-helicity as the peptide traverses the transition path. When we compare this to the free energy, we see a monotonically changing free energy landscape. Contrasting this to S-I for the PE:PG:CL membrane (Fig 7B), we observe a contrasting trend, with initial unfolding, and the abrupt appearance of helicity at point 10, associated with the maxima in the free energy (Fig 4). Importantly, multiple initiation points of helicity (residues K3, L4 and F5 and residues I8, G9 and A10) appear on the peptide. Further folding in the S-I pathway is energetically downhill, with the helical region spanning K3 to L12. S-II (Fig 7C), shows yet another progression, where the initial unfolding is similar to S-I. However, helix formation is now initiated primarily at residues A10, V11, and L12. This folding pathway is energetically favorable, similar to the previous two cases; however, unlike the S-I case, a large energy barrier for the initial folding is not observed, with the differences seemingly arising from the location of helix initiation.

**Figure 7:**
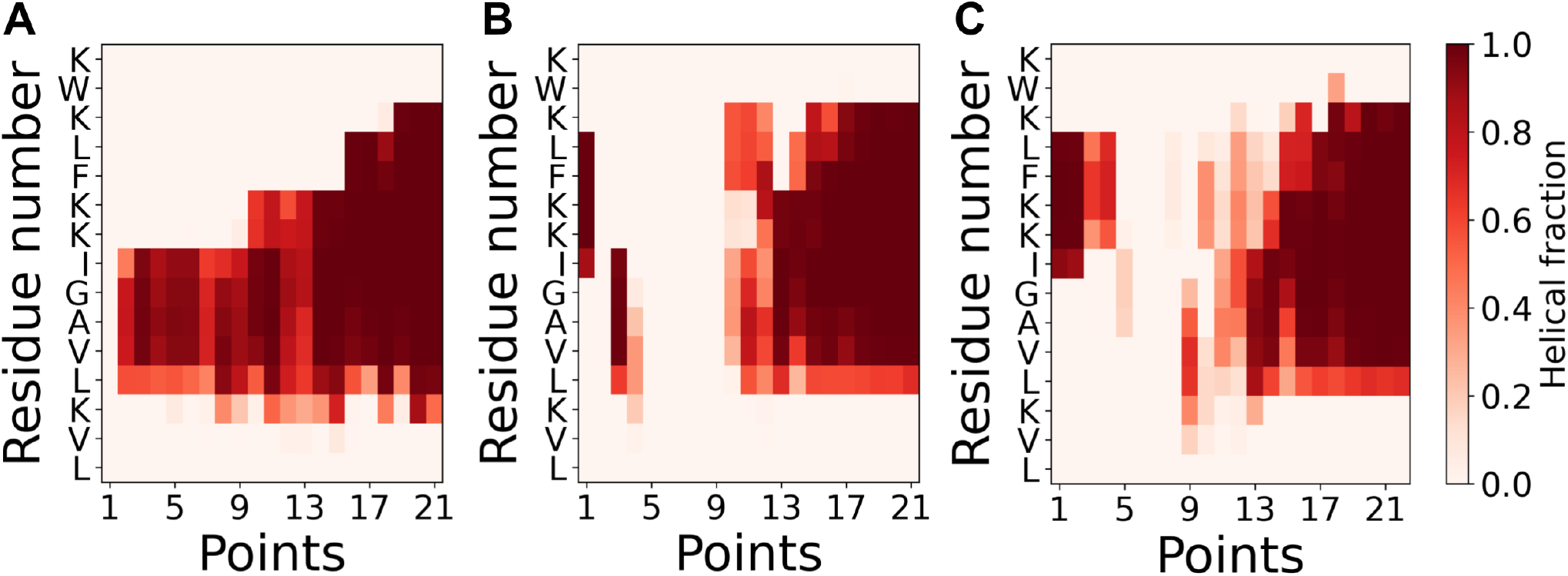
CM15 helicity progression: Fraction of time spent as an *α*-helix for different residues, for the simulation points along the path for (A) PE:PG membrane, (B) Path S-I for PE:PG:CL membrane, (C) Path S-II for PE:PG:CL membrane.

### Cardiolipin alters CM15 hydrogen bonding and lipid contacts along the insertion and folding pathway

We next analyzed hydration, intra-peptide hydrogen bonds (H-bonds), and lipid contacts of the peptide along the path plotted as a function of the points along the converged string as illustrated in Figure 8. The different states are marked for comparison with the string evolution and the free energy profiles. Hydration is a measure of the peptide-water hydrogen bonds (see Methods). The *M*_*bu*_ state is associated with a drop in hydration, and the extent of decrease is similar for all three cases (Figure 8A,B, and C). As folding progresses, the hydration decreases and the H-bonds increase for all cases; however, distinct differences are observed between the PE:PG membrane (Figure 8A) and the PE:PG:CL membrane (Figure 8 B and C). Upon transitioning from the *M*_*bu*_ to the *M*_*iu*_ states, a marked decrease in both hydration and H-bonds is observed for the S-I and S-II string evolutions with CL. This is in contrast to the more gradual and continuous changes observed for the PE:PG membranes. We connect this with the formation of the completely unfolded state for the S-I and S-II paths, in contrast to the partially folded intermediate in the PE:PG membrane. A maxima in the free energy (Figure 4B) for the S-I path coincides with a sharp drop in hydration at the *M*_*iu*_ state (Figure 8 B) associated with the deeper insertion of the peptide for this path. Transitioning to the *M*_*if*_ state occurs with a further decrease in hydration and a monotonic increase in the H-bonds for all three cases. Since CM15 is positively charged, we profile the contacts (Equation 3) with lipids to elicit variations in the presence of CL (Figure 8) last row). The PG lipid contacts increase for the PE:PG case upon membrane insertion and folding. However, greater variation in the contacts is observed for the PE:PG:CL membranes, with CL contributing to an unusually large number of contacts despite the low concentration (5%) in the membrane. This provides the first direct molecular evidence of the modulatory role played by CL in the CM15 folding free energy. Interestingly, for the S-I pathway, where the maximum in the free energy is observed during the formation of the *M*_*iu*_ state, CL shows a similar number of contacts when compared with the PG. For the S-II pathway, where the free energy maximum is absent, CL contacts are distinctly lower. To further dissect the role of CL, we evaluate lipid contacts from unrestrained MD simulations. These results are discussed next.

**Figure 8:**
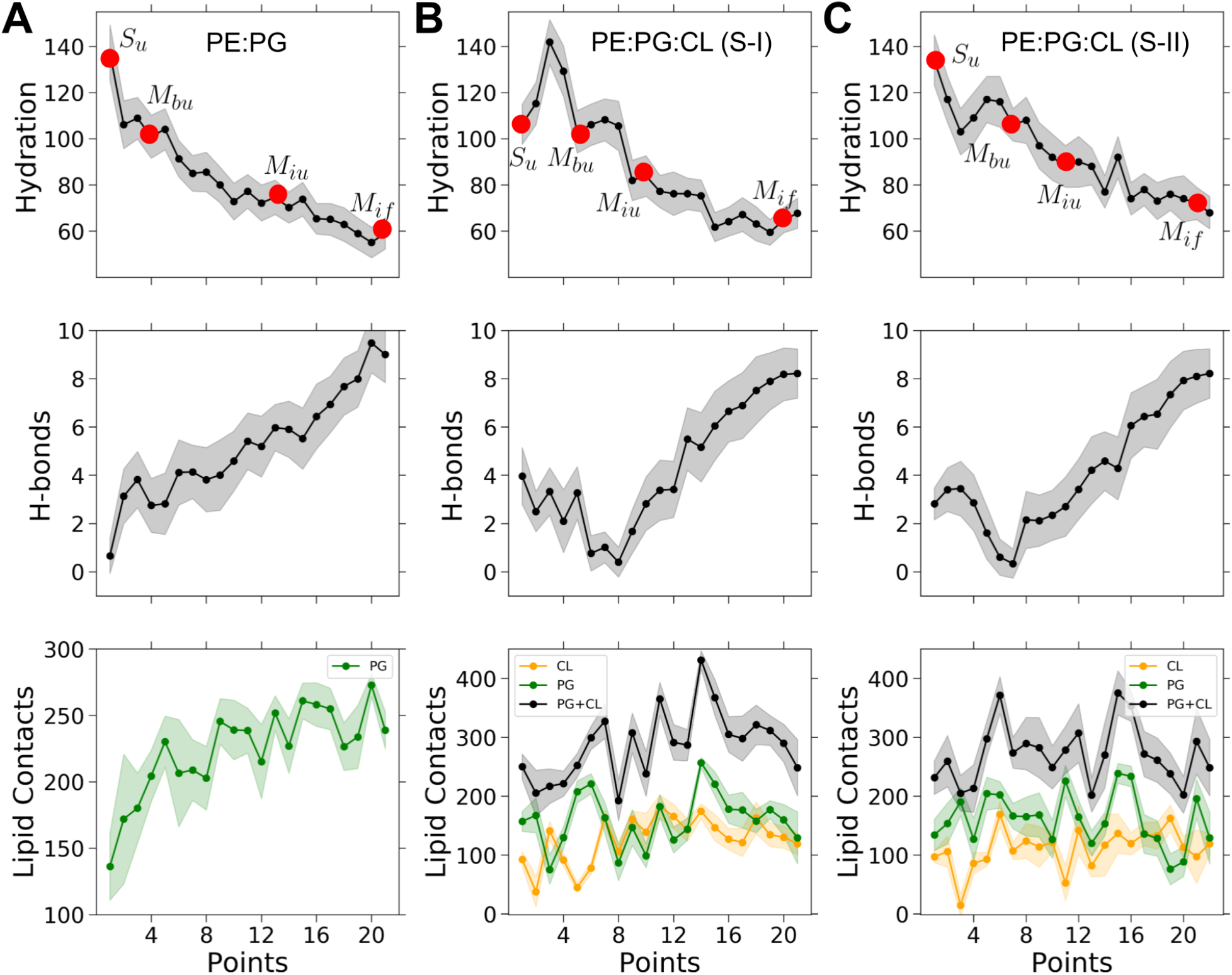
Evolution of various properties for CM15 along the transition path: (A) PE:PG membrane, (B) PE:PG:CL membrane (S-I), (C) PE:PG:CL membrane (S-II). For each case, the hydration, H-bonds, and lipid contacts are illustrated along the column. Data is shown at different points along the string. Hydration represents water molecules present in the hydration shell (0.35 nm) around the peptide, H-bonds are the intra-peptide hydrogen bonds along the path, and contacts are between the peptide and different lipids (see methods).

### CM15 preferentially interacts with cardiolipin in membrane-bound states

The contacts of the different lipid components with the peptide are analyzed to discern preferential interactions between the lipid molecules and understand how low concentrations of CL in the membrane could drastically influence the peptide free-energy landscapes for membrane binding and subsequent folding. Lipid contacts with CM15 residues are normalized by the number of the specific lipids present in the membrane. Figure 9A illustrates the lipid contacts analyzed from the unbiased 4 *μ*s simulation trajectory (same as Figure 2), where the peptide resides predominantly in the M_*bu*_, unfolded state. In contrast, Figure 9B is obtained for a fully folded peptide state, (M_*if*_ ) initially placed beneath the headgroups (see Figure S11) For the M_*bu*_ state, despite the small fraction, CL lipids dominate the contact profiles across all the residues in the vicinity of the N terminus, rich in positively charged lysine (K) residues, showing the largest number of contacts. A similar situation is observed for the M_*if*_ state (Figure 9B). This finding is consistent with the observations of McCammon and co-workers [34, 62] where the tryptophan residue located toward the N-terminus was found to play a key role in membrane interactions for melittin, cecroporin A, as well as CM15.

**Figure 9:**
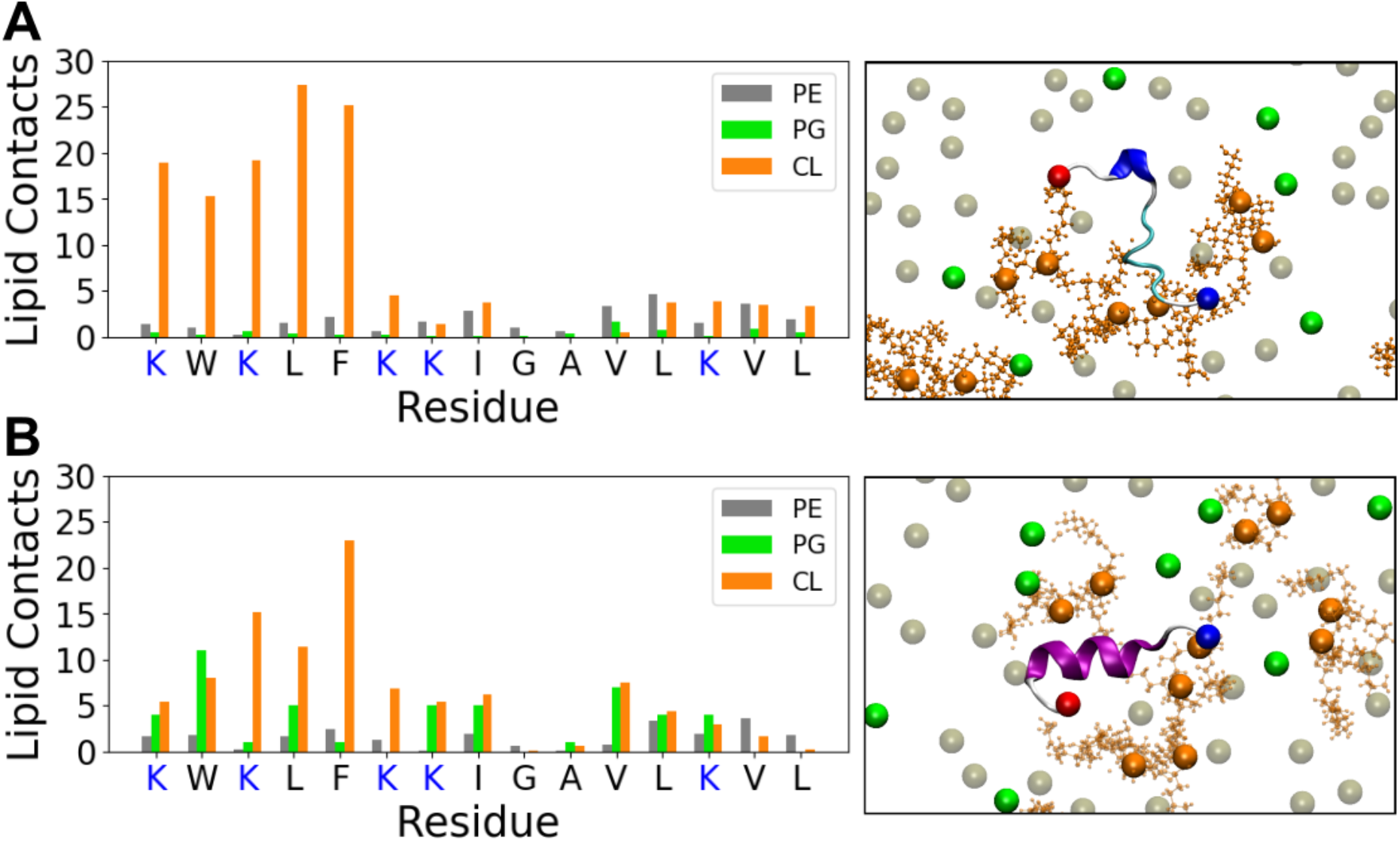
Lipid contact analysis in PE:PG:CL membrane. Lipid contacts with each residue of the peptide from the final 500 ns of the unrestrained simulations for (A) membrane-bound unfolded (*M*_*bu*_) peptide (Figure 2) and (B) inserted membrane-parallel (*M*_*if*_ ) folded peptide (Figure S11). Positively charged residues, K are depicted in blue. The corresponding final topview snapshots of the peptide and membrane are also illustrated. Phospholipid headgroups are depicted as grey-PE, green-PG, orange-CL. The *α*-helical sections of the peptide are highlighted in purple. The N and C-termini are shown in blue and red respectively.

The corresponding snapshots for both states illustrate a strong correlation of CL molecules with the peptide. In comparison with contacts for the PE:PG membrane (Figure S12), a distinct preference for CL binding is observed in the PE:PG:CL membrane. These results suggest that the binding preference for CL is not purely electrostatic and could be connected with membrane deformations associated with the four-tailed CL lipids. With the low CL content in the PE:PG:CL membranes, it is likely that the CL composition around the bound peptide is an additional slow variable resulting in compositional heterogeneity during peptide binding. We see this playing out with contacts analyzed along the string (Figure 8). These variations suggest that the local lipid composition around the peptide can play a role in modulating the free energy landscape, giving rise to the different pathways captured in part with the S-I and S-II evolutions observed in this study.

### CM15 and cardiolipin co-localize in regions of negative membrane curvature

Since CL is known to partition in membranes towards regions with negative curvature [63, 64], we performed curvature and thickness calculations for the unrestrained trajectories used in Figure 9 to uncover correlations with peptide-CL binding. For ease of visualization, the peptide was centered in the simulation box. We performed the curvature analysis for the extracellular leaflet over the final 500 ns of the respective simulations and contrast the variations between the *M*_*bu*_ and *M*_*if*_ states in Figure 10A and B, respectively. In both cases, CL is observed to be co-localized with the peptide. The extent of co-localization is enhanced for the fully folded, *M*_*if*_ state (Figure 10B) compared to *M*_*bu*_. Since CL is present in low concentrations, we overlay the 2D densities over the last 500 ns (shown in black) of the CL headgroups on the curvature maps. The location of the peptide residues is averaged in a similar manner (shown in yellow). For both states, a strong correlation between CL and the negative-curvature regions is observed. This correlation is greater for the folded peptide (Figure 10B). CL is also found to be strongly correlated with the negative curvature regions in the absence of the peptide (Figure S13B). The peptide-CL interactions are also evident for both peptide states. To further quantify this co-localization, we evaluated the probability of locating the center-of-mass of the peptide and the center-of-mass of CL with the negative and positive curvature extrema on the membrane. In all cases, the probability of locating either the peptide or CL is higher for the negative curvature extrema (Figures 10 C-F). The extent of co-location of the peptide with the negative curvature region is the greatest for the *M*_*if*_ state as observed in Figure 10D. Based on these MD simulation results, CM15 is found to interact strongly with CL, despite the small fraction of CL in the membrane, resulting in a co-localization of the peptide with CL in negative curvature induced regions. Combined with our free-energy analysis, the presence of CL in low quantities should impede pore formation associated with downstream AMP activity of CM15. To test this hypothesis, vesicle leakage experiments with varying CL content were carried out. We next discuss these results.

**Figure 10:**
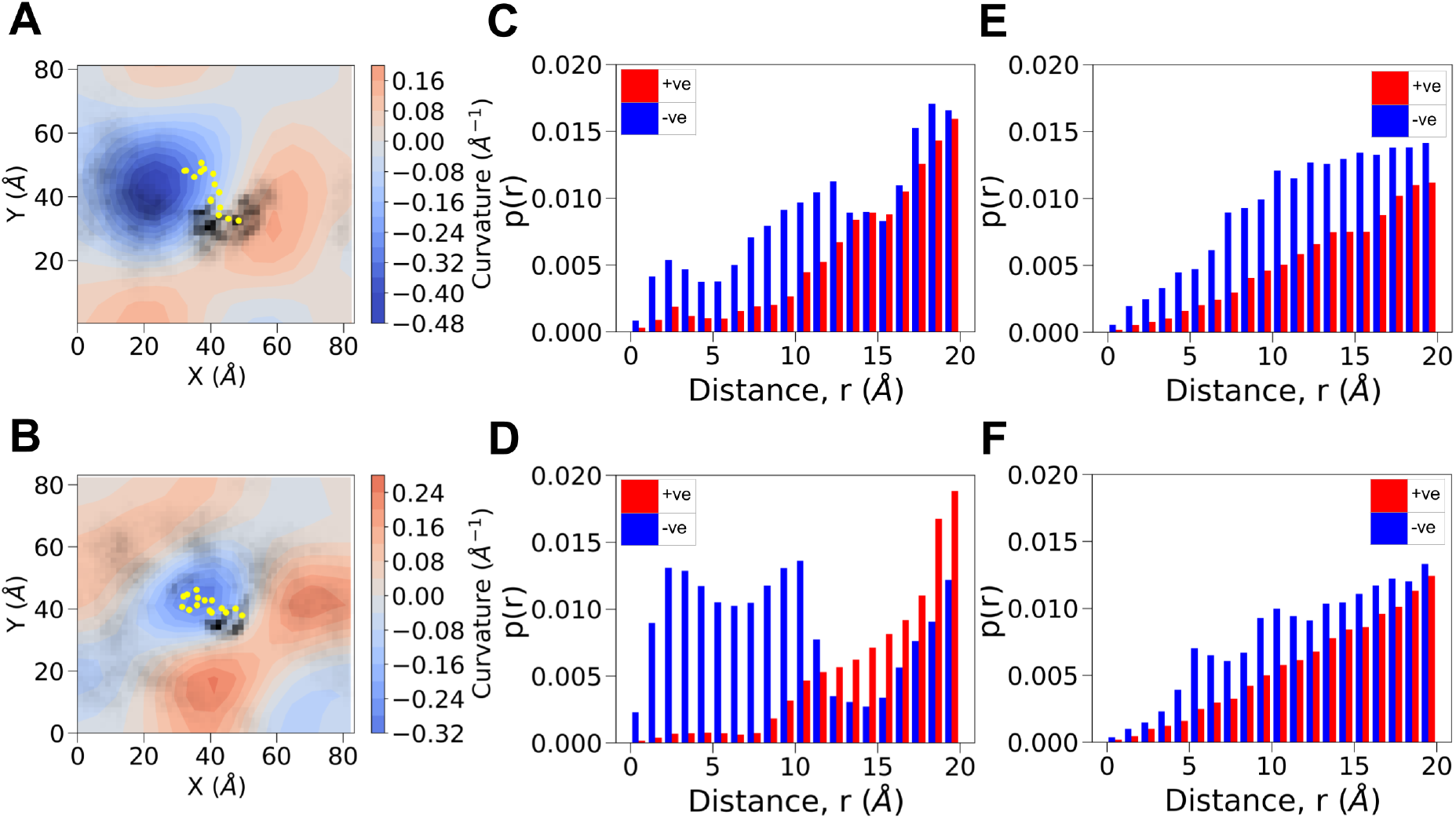
Membrane curvature analysis and CM15-Cardiolipin co-localization. Membrane curvature for the extracellular leaflet averaged over the final 500 ns of the unrestrained simulation trajectories for (A) the membrane-bound unfolded (*M*_*bu*_) peptide configuration (Figure 2), and (B) inserted membrane-parallel (*M*_*if*_ ) folded peptide configuration (Figure S11). The average position of peptide residues is shown in yellow. Additionally, the density of CL lipid in the X-Y plane is plotted in black, overlaid on the curvature plot. Distance from the positive and negative curvature extrema for (C) peptide and (E) CL of the *M*_*bu*_ state. The distance from the positive and negative curvature extrema for (D) peptide and (F) CL for the *M*_*if*_ state.

### Vesicle leakage reveals cardiolipin suppression of CM15 activity

To evaluate the hypothesis that CL interferes with the membrane insertion and refolding processes, we performed dye-release assays using sulforhodamine B-loaded PE:PG:CL vesicles, whose CL content was adjusted between 0 and 20 mol%, thereby covering the range documented for various bacterial species [13, 65]. CM15 was applied at peptide-to-lipid (P:L) ratios spanning 1:100 to 1:1, and membrane leakage was monitored over a 20-minute period. Figure 11A shows the fractional leakage (Equation 5) measured after 20 minutes as a function of P:L ratio for each CL composition. Two distinct regimes are apparent. At low P:L ratios (1:100 and 1:20), the extent of leakage decreases monotonically with increasing CL mole fraction. Specifically, at P:L = 1:20, the fractional leakage decreases from 0.33 in the absence of CL to 0.08 at 20% CL, corresponding to a four-fold reduction. For P:L ratios above 1:10, the fractional leakage plateaus between 0.5 and 0.6 for all membrane compositions, and the dependence on CL inverts, such that CL-containing vesicles exhibit slightly greater leakage than PE:PG vesicles. The transition between these two regimes occurs at P:L ≈ 1:10, where the leakage profiles for all CL contents intersect, although we still observe the leakage reduction at 20% CL. Since CL increases the membrane’s net negative charge, enhanced CM15 binding may contribute to the elevated leakage at P:L ratios exceeding 1:10.

**Figure 11:**
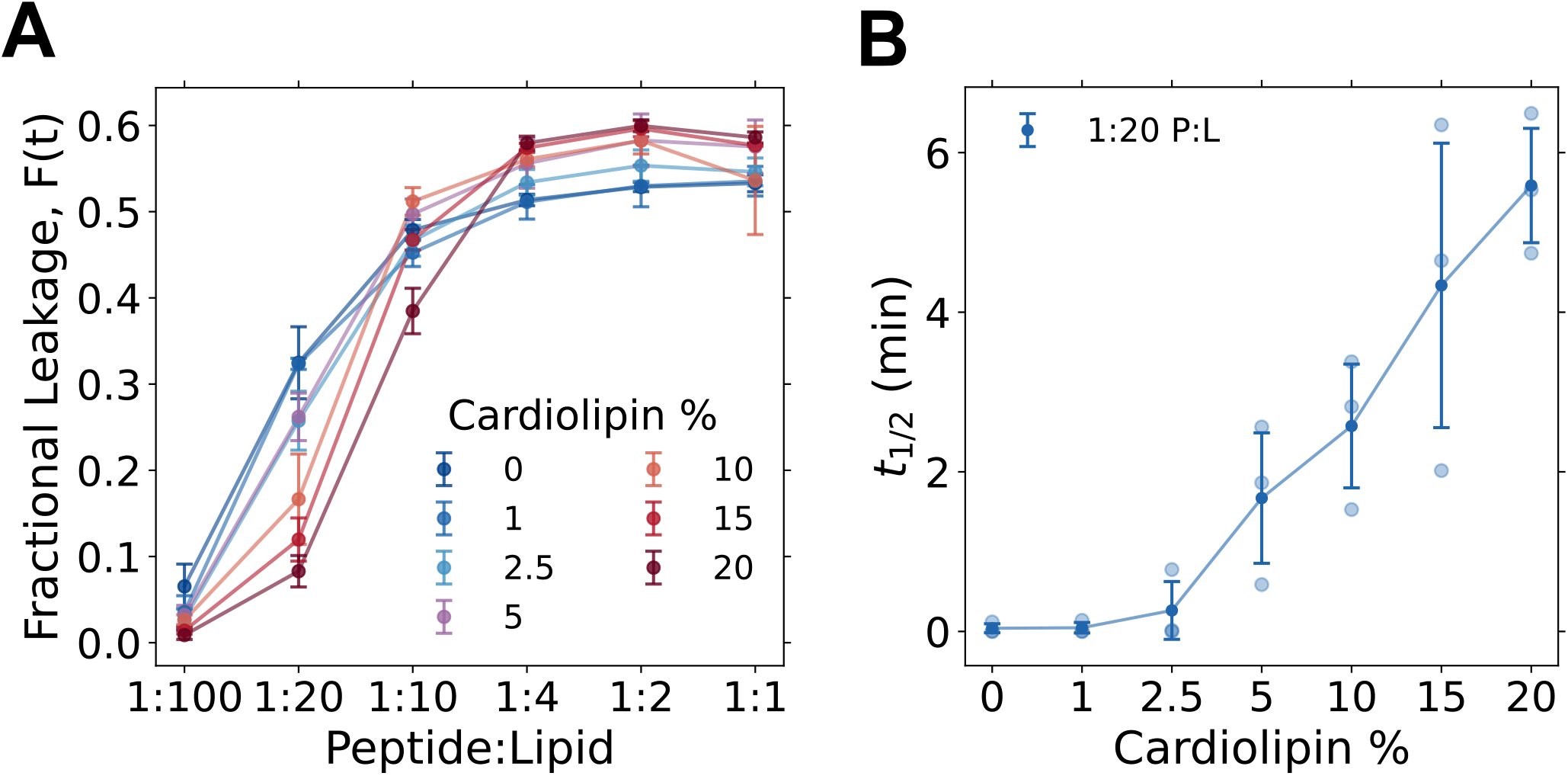
Cardiolipin inhibits CM15-induced vesicle leakage at low peptide-to-lipid ratios: (A) Fractional leakage from sulforhodamine B-loaded vesicles after 20 minutes of CM15 treatment, as a function of peptide-to-lipid ratio, for CL contents from 0 to 20 mol%. (B) Leakage half-time *t*_1/2_ at a peptide-to-lipid ratio of 1:20 as a function of CL content; light symbols denote individual measurements and dark symbols the mean. Half-times at 0 and 1% CL fall below the time resolution of the measurement. Error bars denote standard deviations over three independent preparations, n = 3.

The CL mediated activity suppression is also seen by visualizing the normalized leakage ratios, defined as the fractional leakage of PE:PG:CL vesicles to the fractional leakage of PE:PG vesicles (Figure S14). At lower P:L (below 1:10), the leakage ratio decreases as cardiolipin concentration increases, indicating suppression of activity. For P:L ratios above 1:10, the leakage ratio remains close to or above 1.0, indicating no suppression or mild enhancement. Since a free energy barrier governs the rate at which a state is attained rather than its equilibrium population, we next examined the leakage kinetics directly. Figure 11B shows the leakage half-time, *t*_1/2_, at a P:L ratio of 1:20, the condition under which half-times can be determined with suffcient reliability (see Methods). In the absence of CL, and at 1% CL, leakage is complete within the instrumental dead time, and *t*_1/2_ cannot be resolved. At CL contents ≥ 2.5%, the half-time increases steeply and monotonically, attaining values of ≈ 2.6 minutes at 10% CL and ≈ 5.6 minutes at 20% CL. Thus, over this compositional range, CL slows the approach to the leakage plateau by more than an order of magnitude.

This kinetic signature provides the most direct experimental correspondence with the free energy calculations. The simulations attribute the principal CL-dependent free energy barrier to the insertion step, during which the unfolded peptide is positioned beneath the phospholipid headgroups, with subsequent folding proceeding downhill in free energy. A barrier at this stage slows the rate at which peptides attain the folded, membrane-parallel conformation required for pore assembly and is therefore expected to manifest as a reduction in the leakage rate, rather than solely as a reduction in the overall leakage amplitude. The observed increase in *t*_1/2_ with CL content, together with the decrease in leakage amplitude at low peptide-to-lipid ratios (P:L), is consistent with this mechanistic interpretation. Notably, at 5% CL—the composition used in the free energy calculations and representative of the *E. coli* inner membrane-the half-time is already elevated to approximately 1.7 minutes from an unresolved value in the CL-free membrane, indicating that the effect is substantial at physiologically relevant CL levels.

## DISCUSSION

Using all-atom simulations and an enhanced-sampling path-based method, we provide a detailed mechanism for CM15 insertion and folding in bacterial membranes, with a specific emphasis on the role of CL, a four-tailed negatively charged lipid found in both Gram-positive and Gram-negative bacteria. Vesicle leakage experiments at different peptide-to-lipid ratios provide a link between the folding energetics of a single CM15 peptide and the downstream mechanisms of CM15 driven leakage activity. The string-method free-energy analysis in a two-dimensional collective-variable space illustrates the intimate interplay between membrane insertion and subsequent folding of the peptide below the phospholipid headgroups. Our analysis provides molecular details of the previously proposed membrane-binding and subsequent refolding behavior of CM15 [32, 66]. We show that membrane insertion initially occurs with the peptide in an unfolded state, followed by folding into a fully helical membrane-parallel state. In the PE:PG membrane, unfolded CM15 spontaneously inserts into the membrane and subsequently folds into an *α*-helical state. However, in the presence of CL, the S-I and S-II strings reveal two distinct pathways (Figure 4 and 5). These pathways reflect the inherent ruggedness of the free-energy landscape arising from variations in the local association of CL with the peptide. We attribute these changes to strong CM15-CL interactions and the resulting perturbation of the local lipid environment around the bound peptide. Thus, even at a low concentration, CL substantially modifies the free-energy landscape associated with CM15 insertion and folding. The emergence of two pathways also highlights the multidimensional nature of the underlying free-energy landscape. Delineating the complete landscape would require expanding the collective-variable space to include the local lipid composition around the peptide, specific peptide-lipid contacts, and potentially other slow variables not explicitly considered here [67].

The vesicle leakage experiments reveal two distinct regimes. At low P:L ratios (< 1:10), an increase in CL content, reduces the vesicle leakage. However, at higher P:L ratios, leakage is independent of CM15 concentration, with a marginally enhanced leakage readout with higher CL content. The crossover between the CL dependent behaviors occurs near P:L ∼ 1:10. Our free-energy computations shed light on the initial steps of AMP activity associated with membrane binding and resulting conformational transitions. Pore formation is a collective effect that arises with increased peptide concentration, a combined effect due to bilayer thinning and defect mediated poration. CM15 has previously been proposed to permeabilize membranes through a toroidal pore-like mechanism, where both peptides and lipid headgroups line the aqueous transmembrane pore [68–70].

Although the microscopic pathways of CM15-induced leakage remain elusive [71], pore formation provides a direct link between the single-peptide insertion and folding processes characterized in our simulations, to the vesicle leakage readouts in the experiments.

At the lower CM15 concentrations (< 1:10, P:L ratios), we posit that toroidal pore formation with increasing CL content is hindered, since packing of the negative curvature forming CL molecule into the positively curved regions of the pore would be energetically unfavorable. This mitigating influence of CL on pore-formation free energies has recently been studied by Rocha-Roa et al. [27], supporting this argument. The presence of CL would hinder a mechanism that requires lipid-associated peptides to reorient to form the toroidal pore complex. MD simulations with the AMP, aurein 1.2 also reveal the mitigative influence of CL on pore formation [28]. Strong binding of CM15 to CL, as seen in our MD simulations, is likely to result in pinning of the peptide to the lipid headgroup region to further impede pore formation, with the resulting inhibitory effect on vesicle leakage with increasing CL content. This effect is observed for CM15 concentrations at low P:L ratios.

At higher P:L ratios, the accumulation of CM15 at the membrane interface is expected to increase the importance of collective peptide-induced membrane perturbations. In this regime, a high surface density of CM15 can promote cooperative membrane thinning, defect formation, and poration. The experimentally observed leakage plateau is therefore interpreted as a saturation of membrane permeabilization, as the amount of CM15 binding to the membrane is maximized. Similar saturation effects akin to Langmuir isotherms have been reported in vesicle leakage and peptide-vesicle binding studies [71, 72]. We note, however, that the enhancement in leakage in this regime is small, indicating that the increase in the total number of pores formed is not a strong function of CL concentration at these high P:L ratios. Finally, we note that we do not observe any signatures of detergency mediated vesicle disruption activity at the highest CM15 concentration considered in this study.

## CONCLUSION

Although AMP activity has been extensively investigated, the free-energy changes associated with membrane binding, insertion, and folding, as well as the differences arising from compositional variations in bacterial cells, have not been reported previously. Our study reveals the balance between peptide concentration, CL content, and ensuing pore formation. The free energy barrier for the transition of CM15 from an unfolded state to a membrane folded state is enhanced in the presence of CL. Our study shows that this effect is strong, even at low CL concentrations of 5% driven primarily by the electrostatic interactions of the positively charged CM15 peptide and the anionic CL. The resulting landscape is rugged, as revealed by the emergence of different minimum free energy paths. Vesicle leakage experiments reveal a distinct crossover from a CL inhibited regime at low peptide concentrations to a saturation regime with a weakly enhanced leakage activity as CL concentration is increased. Our study has direct implications for the design of AMPs for antibacterial activity, illustrating the putative role of CL in pore formation. Recent lipidomics data [14] for Gram-positive strains reveal CL content ranging from 5 - 75% with higher CL content associated with extremophiles surviving in high saline environments. We expect our results to apply across a wide class of amphipathic AMPs, providing molecular insights to assist in the search for strain-specific antimicrobial molecules.

## DATA AVAILABILITY

Information required to reproduce the simulation data using the open-source GROMACS and PLUMED is provided in the manuscript or included in the Supplementary Information.

## AUTHOR CONTRIBUTIONS

KGA,SP and RR designed the research and formulated the problem. NMB and AK carried out simulations and conducted the free energy analysis. NMB analyzed the trajectories. AU carried out the vesicle leakage experiments. All authors were involved in writing initial drafts and editing the article.

## ACKNOWLEDGMENTS

We acknowledge the Supercomputer Education and Research Center (SERC) computing facility; Thematic Unit of Excellence on Computational Materials Science (TUE-CMS), a Department of Science and Technology (DST)-supported computing facility at the Indian Institute of Science Bangalore; and the National Supercomputing Mission, India, for funding used in this work.

## Notes

### Competing Interest Statement

The authors have declared no competing interest.

