## Supplementary Information for "Cardiolipin regulates free energy of folding and vesicle leakage of CM15 antimicrobial peptides in bacterial inner membranes"

Table S1: CM15 and bacterial membrane simulation details performed in NPT ensemble at 0.15 M NaCl salt concentration.

| Membrane composition | Simulation time (ns) | Peptide initial state | Peptide initial location | Peptide final state | Peptide final location |
|---|---|---|---|---|---|
| Aqueous | 600 | Folded | Bulk | Unfolded | Bulk |
| | 1000 | Folded | Bulk | Unfolded | Bulk |
| | 1000 | Folded | Bulk | Unfolded | Bulk |
| PE:PG (150:40) | 4000 | Unfolded | Extracellular space | Folded | Below headgroup plane |
| | 1000 | Unfolded | Extracellular space | Partially folded | Below headgroup plane |
| | 1000 | Folded | Extracellular space | Folded | Below heagroup plane |
| | 1000 | Folded | Below headgroup plane | Folded | Below headgroup plane |
| PE:PG:CL (150:40:10) | 4000 | Unfolded | Extracellular space | Unfolded | Headgroup plane |
| | 1000 | Folded | Extracellular space | Unfolded | Headgroup plane |
| | 1000 | Unfolded | Extracellular space | Unfolded | Headgroup plane |
| | 2000 | Folded | Below headgroup plane | Folded | Below headgroup plane |

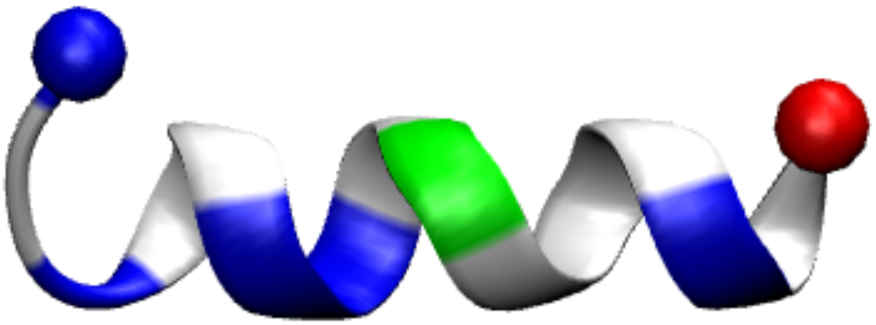

Figure S1: **CM15 crystal structure:** The N and C-termini are represented by spheres with blue and red colour respectively. The positively charged residues are shaded blue, polar residues are shaded green and non-polar residues are uncoloured.

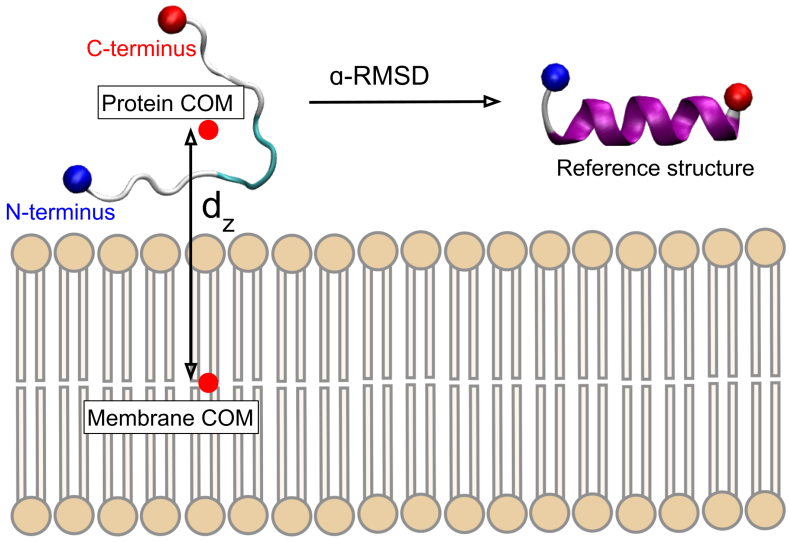


Figure S2: **Schematic for evaluating the considered collective variables (CVs):** $\alpha$−RMSD is calculated in comparison to a reference PDB structure of the peptide. $d_z$ is the $z$-component of the vector between the protein centre-of-mass and membrane centre-of-mass.

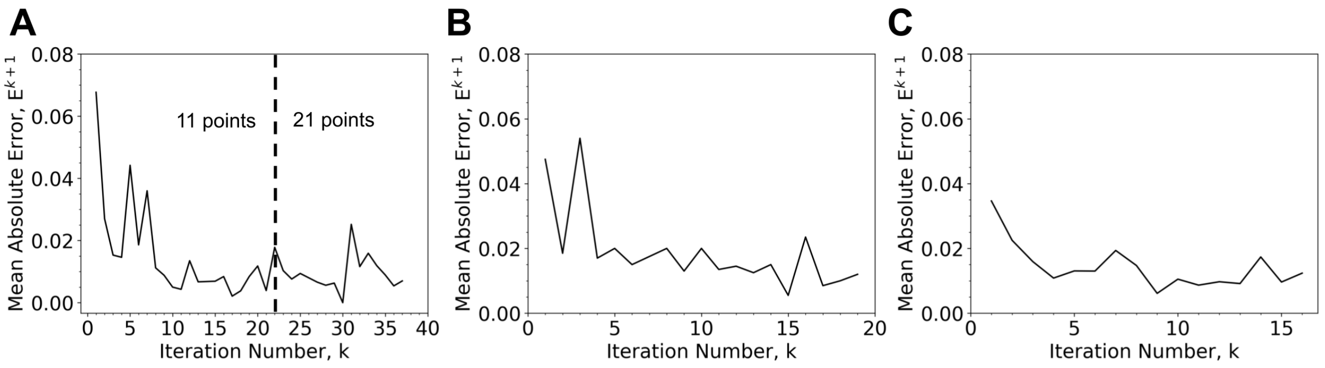


Figure S3: **String convergence.** Mean absolute error, $E^{k+1}$ (see Eq. 2 in the main manuscript) for each iteration of string convergence for A) PE:PG membrane, B) S-I for PE:PG:CL membrane C) S-II for PE:PG:CL membrane

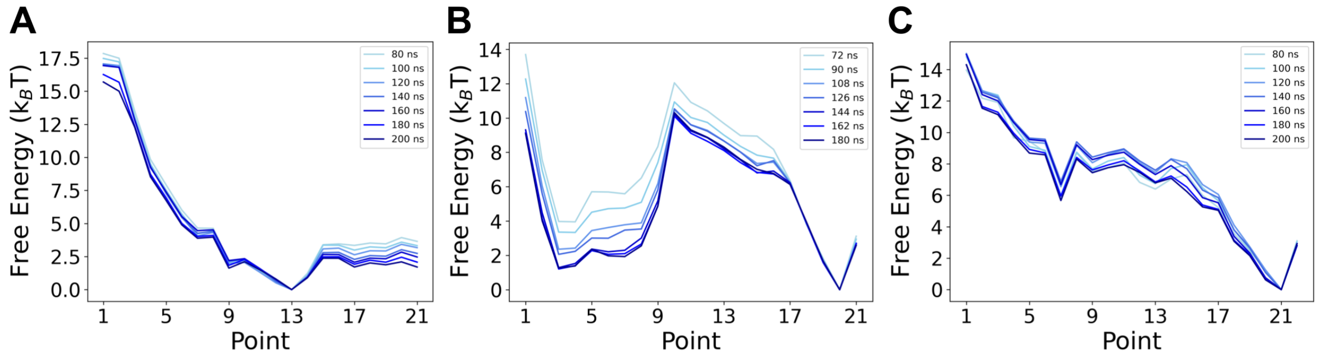


Figure S4: **Convergence of free energy from WHAM.** Free energy along the path derived for A) PE:PG membrane starting at 80 ns per simulation window and recalculated with subsequent additional 20 ns up to 200 ns per simulation window, B) S-I for PE:PG:CL membrane starting at 72 ns per simulation window and recalculated with subsequent additional 18 ns up to 180 ns per simulation window and C) S-II for PE:PG:CL membrane starting at 80 ns per simulation window and recalculated with subsequent additional 20 ns up to 200 ns per simulation window.

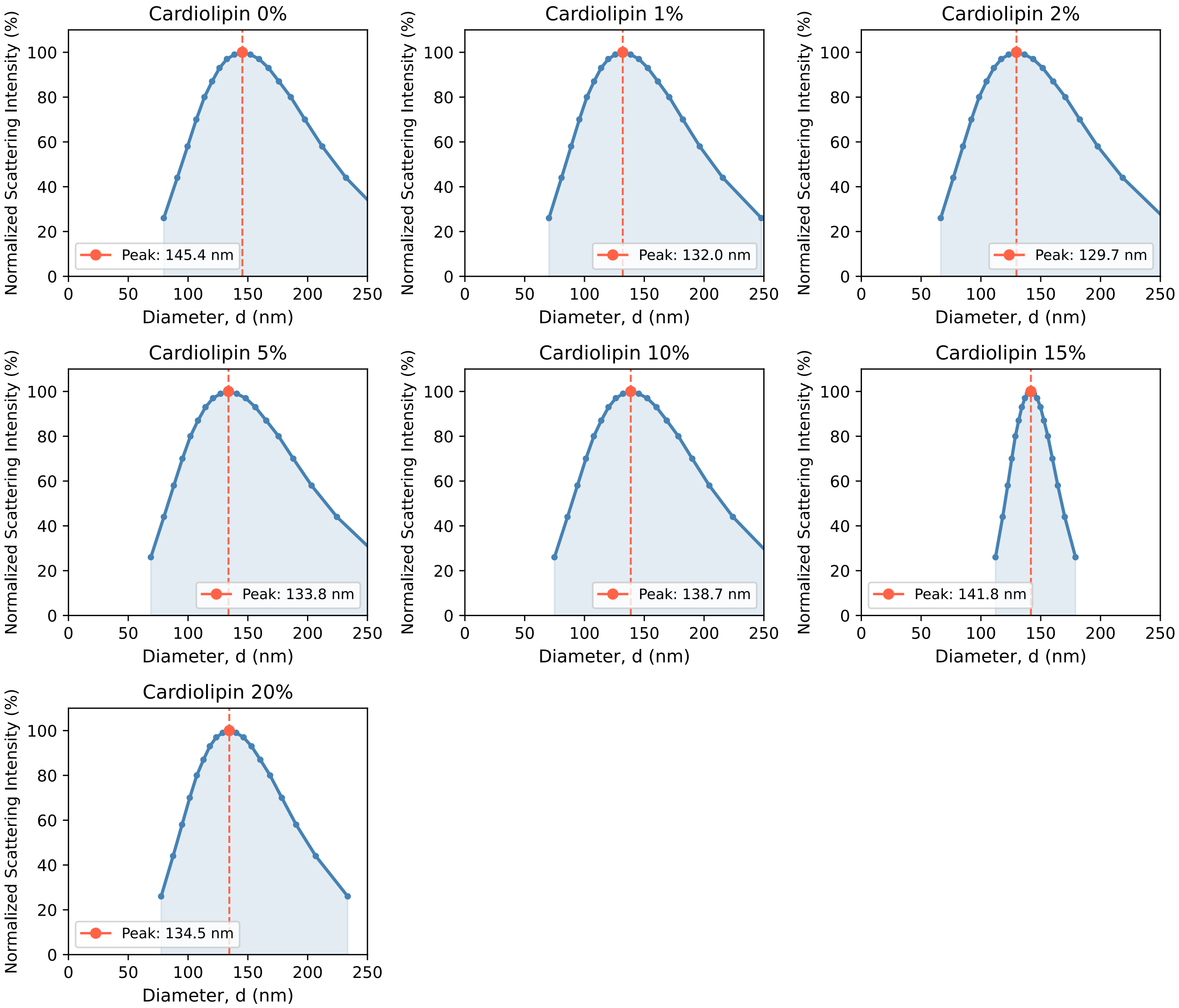


Figure S5: **Vesicle size estimation using Dynamic Light Scattering:** The vesicle size of SRB-loaded vesicles was determined using DLS to verify the extrusion by 0.1 $\mu$m membranes. The peak diameter indicates vesicles between 129 - 145 nm diameter. The vesicles were diluted to 30 $\mu$M concentration before measurement. The distribution shown is an average of 5 repeats.

Table S2: **Vesicle composition in mole fractions for dye-loaded vesicle leakage.**

| % Cardiolipin | DOPE | DOPG | Cardiolipin |
|---|---|---|---|
| 0 | 0.75 | 0.25 | 0.00 |
| 1 | 0.7425 | 0.2475 | 0.01 |
| 2.5 | 0.7312 | 0.2478 | 0.025 |
| 5 | 0.7125 | 0.2375 | 0.05 |
| 10 | 0.675 | 0.225 | 0.10 |
| 15 | 0.6375 | 0.2125 | 0.15 |
| 20 | 0.6 | 0.2 | 0.2 |

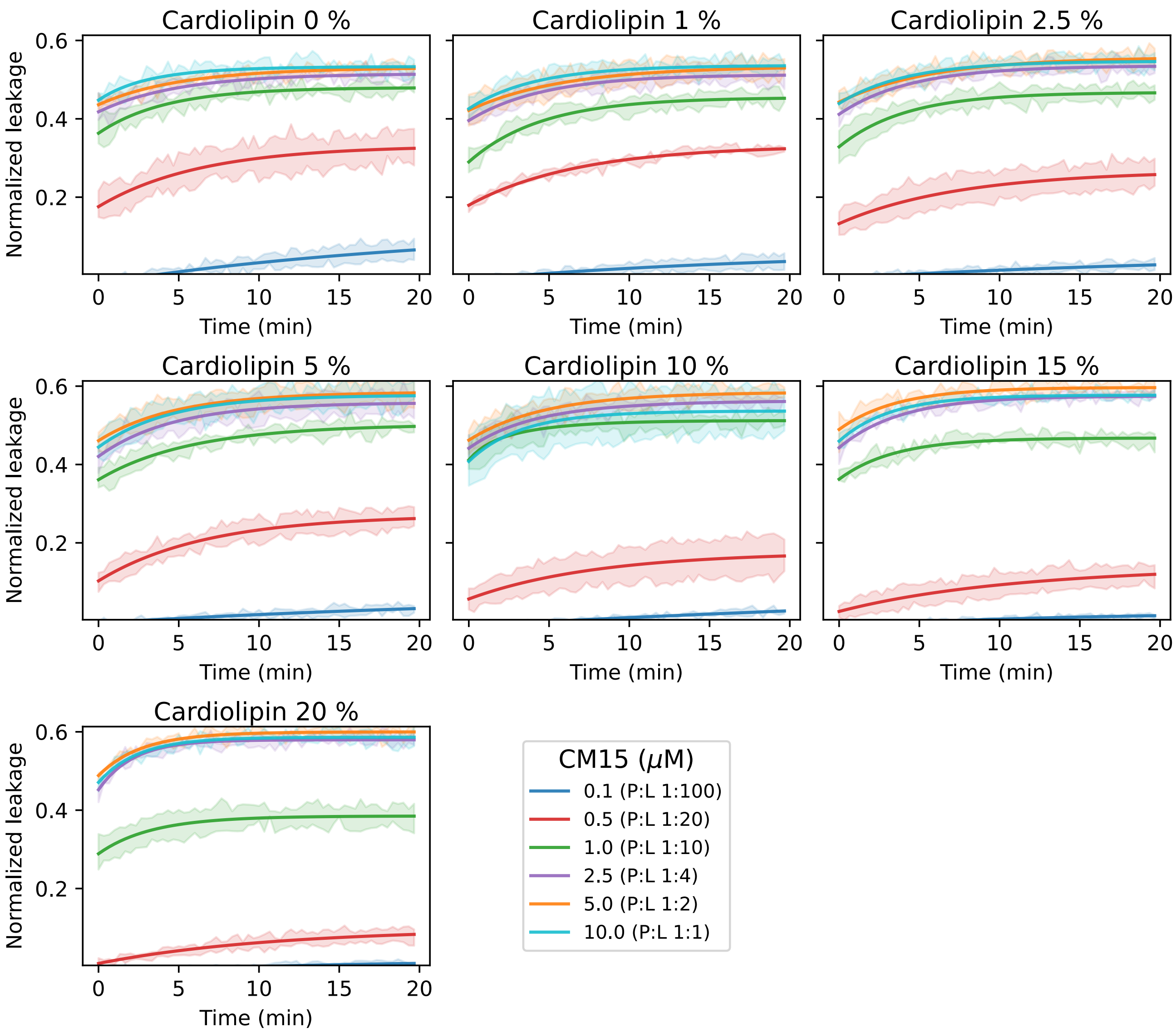


Figure S6: **SRB-loaded vesicle leakage by CM15:** Time traces of vesicle leakage by different CM15 concentrations (0.1, 0.5, 1, 2.5, 5 and 10 $\mu$M, corresponding peptide-to-lipid (P:L) ratios given in bracket) for vesicles having increasing cardiolipin concentrations as given in Table S2. The dark line represents the average value. The semi-transparent region is the standard deviation of the raw data resulting from three replicates. The initial kinetics of the vesicle leakage are not captured due to instrument delay.

Table S3: CM15 concentration corresponding to each peptide-to-lipid (P:L) ratio.

| CM15 ($\mu$M) | P:L ratio |
|---|---|
| 0.1 | 1:100 |
| 0.5 | 1:20 |
| 1.0 | 1:10 |
| 2.5 | 1:4 |
| 5.0 | 1:2 |
| 10.0 | 1:1 |

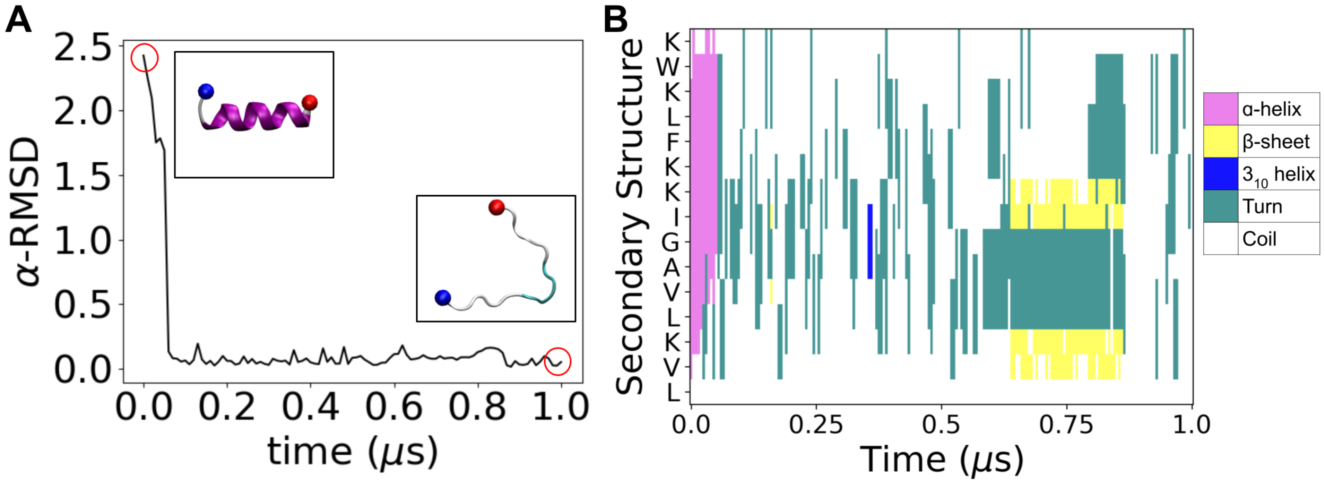


Figure S7: **CM15 aqueous simulation:** CM15 is simulated for 1 $\mu$s in aqueous environment at physiological salt concentration. (A) $\alpha$-RMSD evolution for the peptide over time. The initial and final peptide structure is shown in the inset. The N-terminus is represented by the blue bead, while the C-terminus is represented by the red bead. (B) Residue-wise secondary structure of the peptide over time.

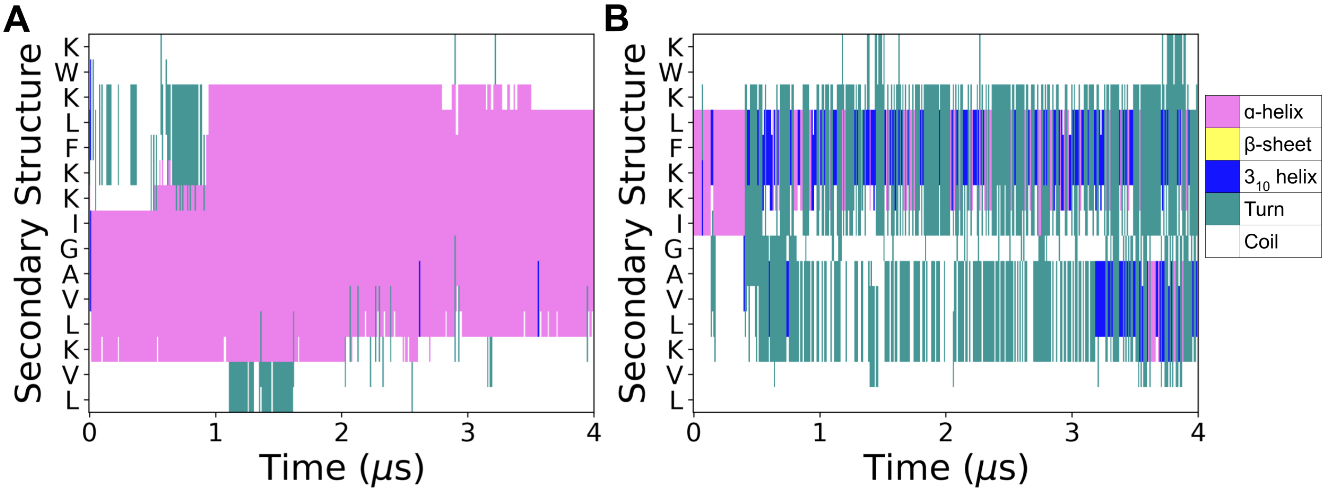


Figure S8: **STRIDE analysis for 4 $\mu$s unbiased peptide-membrane simulations:** Residue-wise secondary structure over time for peptide in (A) PE:PG membrane and (B) PE:PG:CL membrane.

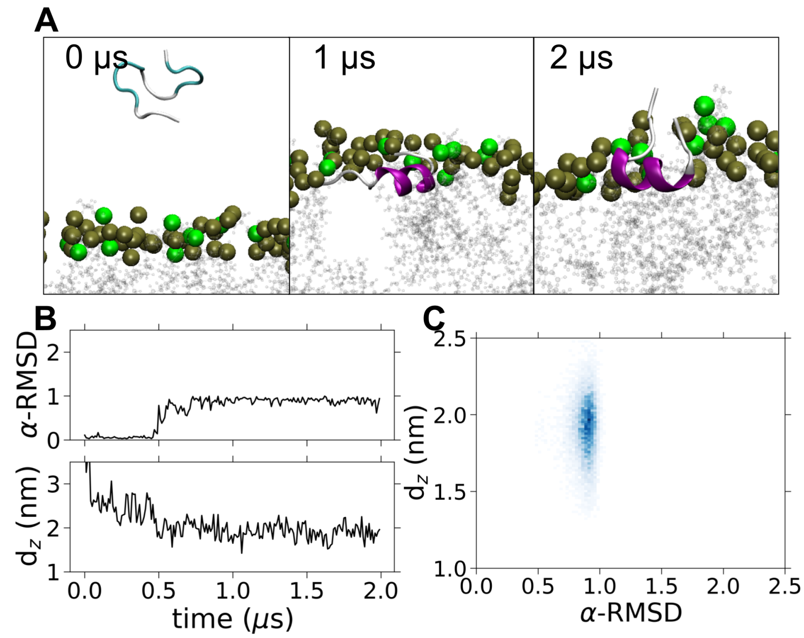


Figure S9: **CM15 and PE:PG membrane simulations.** (A) Snapshots illustrating the orientation and secondary structure adopted by CM15 at different stages of a 2 $\mu$s all-atom MD simulation. (B) The time evolution of the $\alpha$-RMSD and $d_z$. (C) 2D histogram ($\alpha$-RMSD, $d_z$) of the different states sampled across the final 1 $\mu$s of the 2 $\mu$s simulation.

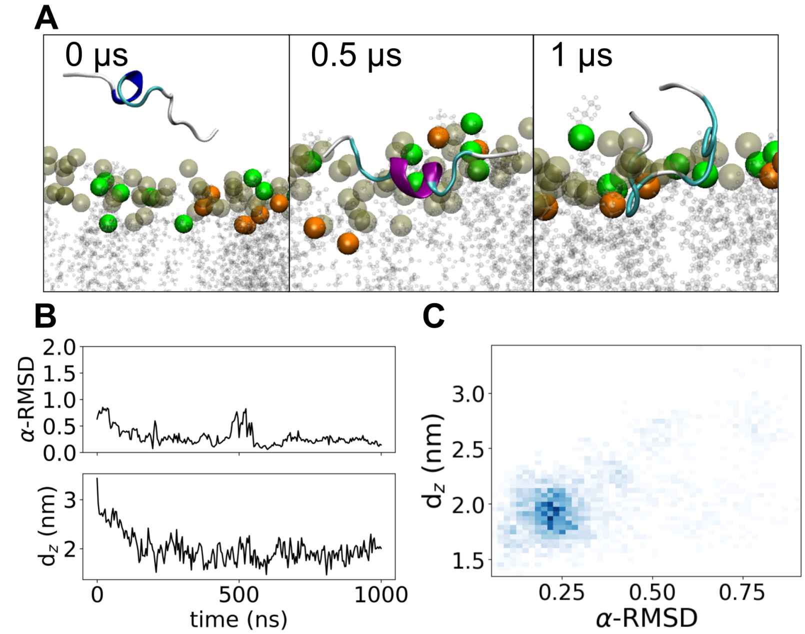


Figure S10: **CM15 and PE:PG:CL repeat membrane simulations.** (A) Snapshots illustrating the orientation and secondary structure adopted by CM15 at different stages of a 1 $\mu$s all-atom MD simulation. (B) The time evolution of the $\alpha$-RMSD and $d_z$. (C) 2D histogram ($\alpha$-RMSD, $d_z$) of the different states sampled across the 1 $\mu$s simulation.

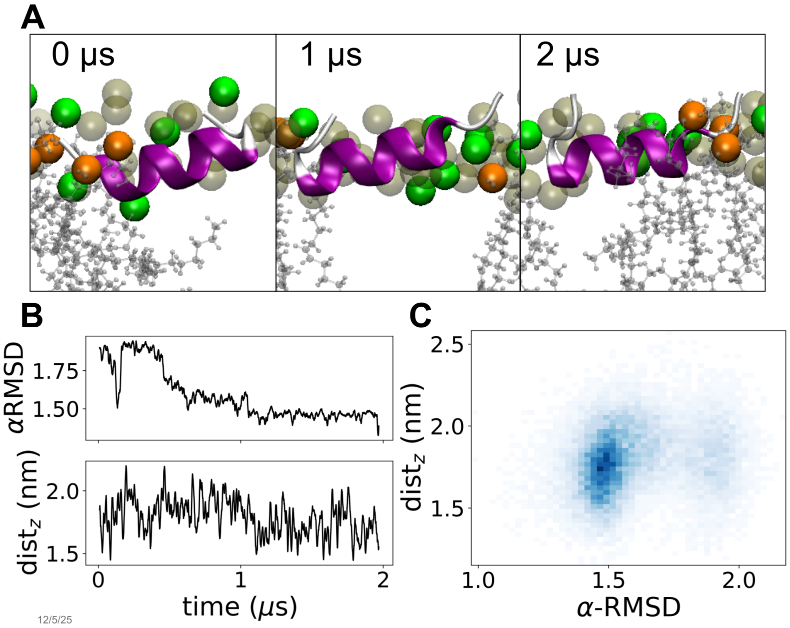


Figure S11: **CM15 and PE:PG:CL membrane simulations.** Simulating the peptide with a membrane-inserted folded ($M_{if}$) initial state. (A) Snapshots illustrating the orientation and secondary structure adopted by CM15 at different stages of a 2 $\mu$s all-atom MD simulation. N-terminus is shown with red and C-terminus is shown with blue sphere. (B) The time evolution of the $\alpha$-RMSD and $d_z$. (C) 2D histogram ($\alpha$-RMSD, $d_z$) of the different states sampled across the final 1 $\mu$s of the 2 $\mu$s simulation.

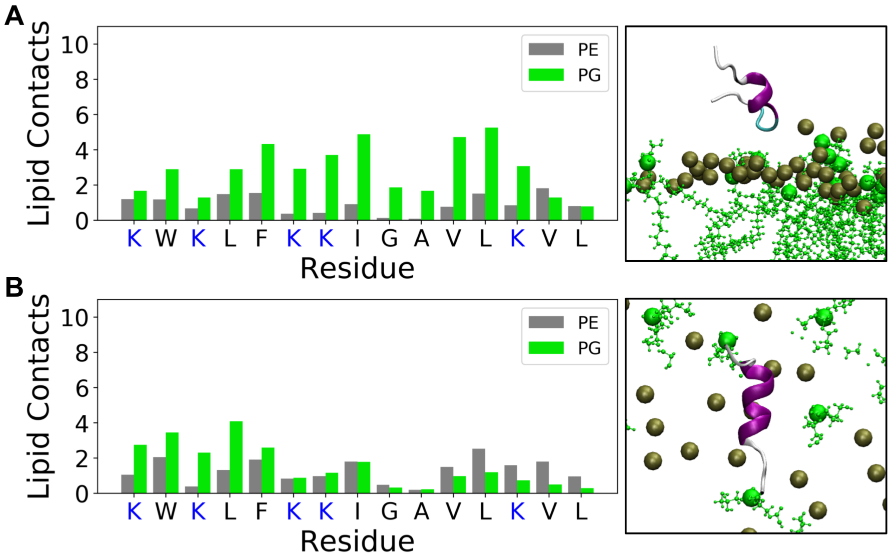


Figure S12: **Lipid occupancy analysis:** Lipid contacts with each residue of the peptide from the final 500 ns of the unbiased simulations for (A) surface unfolded ($S_u$) peptide, and (B) inserted membrane-parallel ($M_{if}$) folded peptide. Positively charged residues are highlighted in blue. The final snapshot of the peptide in the membrane is shown below for MB and MP states, where green colour denotes DOPG molecules. The phosphorus atoms are represented by spheres with colours grey-PE, green-PG. The $\alpha$-helix sections of the peptide are highlighted in purple.

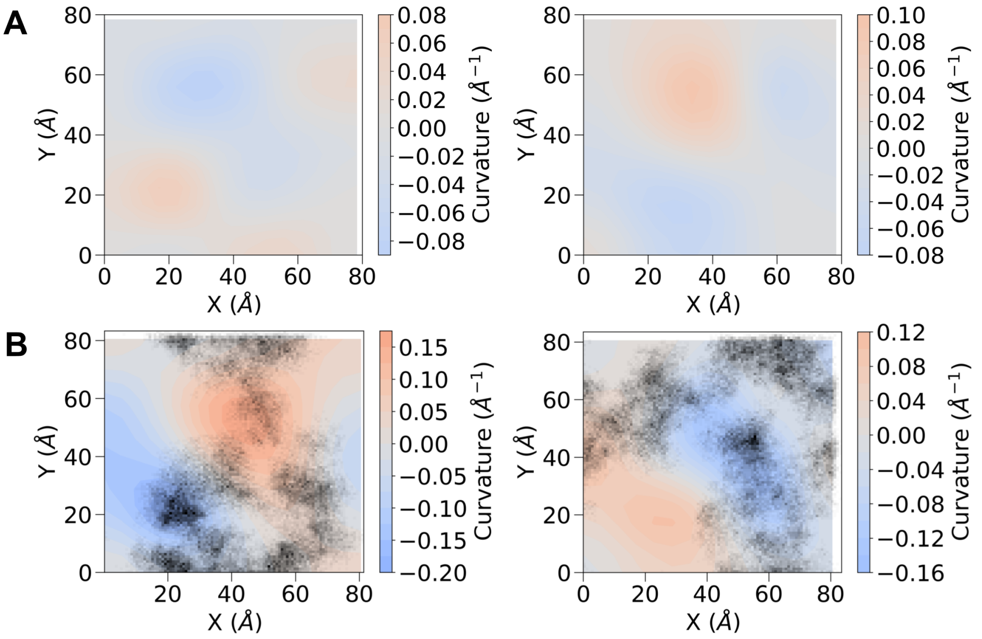


Figure S13: **Curvature analysis for unbiased bare membrane simulations:** Mean curvature for the two leaflets of a bare membrane system for the (A) PE:PG membrane, and (B) PE:PG:CL membrane, with the density of CL lipid in the X-Y plane plotted in black, overlaid on the curvature plot.

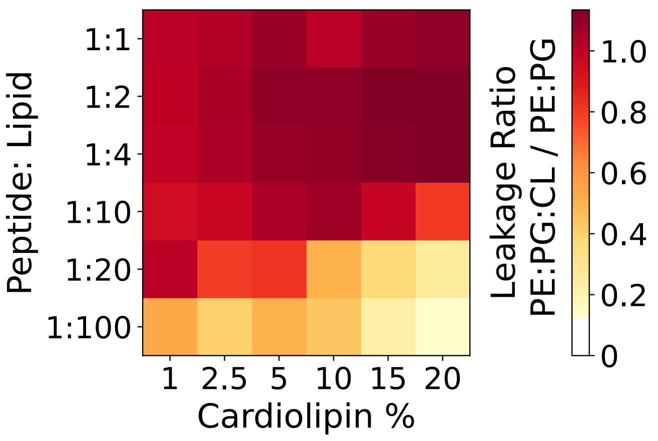


Figure S14: **Leakage ratio heatmap.** Leakage ratio, defined as the fractional leakage of PE:PG:CL vesicles normalised to PE:PG vesicles at the same peptide-to-lipid ratio.
